# Purging of highly deleterious alleles through an extreme bottleneck

**DOI:** 10.1101/2024.07.23.602943

**Authors:** Oliver P. Stuart, Rohan Cleave, Kate Pearce, Michael J.L. Magrath, Alexander S. Mikheyev

## Abstract

Transitions to captivity often produce population bottlenecks. On one hand, bottlenecks increase inbreeding and decrease effective population size, thus increasing extinction risk. On the other, elevated homozygosity associated with extreme bottlenecks may purge deleterious alleles. Previous studies of purging in captive breeding programs have focused on phenotypic measurements and may be confounded with environmental effects. We test the ability of natural selection to purge deleterious alleles following an extreme population bottleneck by analysing patterns of genetic diversity in wild and captive-bred populations of the Lord Howe Island stick insect (*Dryococelus australis*). *Dryococelus australis* has been bred in captivity for two decades, having passed through an extreme bottleneck – only two mating pairs with few new additions since then. The magnitude of the bottleneck, together with the fact that each female lays hundreds of eggs most of which are not recruited, set up nearly ideal conditions for the purging of deleterious alleles. As expected, captive-bred individuals had a greater number of long runs-of-homozygosity compared to wild individuals, implying strong inbreeding in captivity which would facilitate purging in the homozygous regions. Stop-codon and frameshift alleles were preferentially depleted in captivity compared to other alleles, in coding and non-coding regions. The more deleterious an allele was predicted to be, the more likely it was found outside of runs-of-homozygosity, implying that inbreeding events facilitate the expression and thus removal of deleterious alleles, even after such an extreme bottleneck and under the benign conditions of captivity. These data show that extreme inbreeding has likely decreased the deleterious mutation load of the stick insects, though loss of genetic diversity may make future adaptation more difficult.

## Introduction

Captive breeding programs are the only available conservation management action for some severely threatened species. They are likely to become important for more species as human populations continue to grow in the 21st century and progressively more land is cleared for agriculture and other purposes (Soulé et al., 1986). However, the degeneration of fitness in captivity and the divergence of wild and captive phenotypes over time is well known (Crates et al. 2023). Some of this phenotypic change can be owed to the genetic consequences of the demography of captive breeding programs.

Many captive breeding programs begin with a bottleneck event which will increase genome-wide homozygosity at the individual and the population levels (Nei et al. 1975; Chakraborty and Nei 1977). A bottleneck is a sampling process which carries only a fraction of a population’s ancestral diversity into subsequent generations. This sampling process may be stochastic, and so deleterious alleles will be carried through the bottleneck and subsequently exposed in homozygosis. The result is the unmasking of previously hidden recessive deleterious alleles, converting a “masked” genetic load to a “realized” genetic load (Bertorelle et al., 2022) which can be observed empirically in threatened populations by an increase of deleterious homozygous genotypes (Liu et al., 2020; Khan et al., 2021; Mooney et al., 2022; Mochales-Riaño et al., 2023). Additionally, many captive breeding programs have small populations compared to wild populations, amplifying the effect of drift in addition to the effect of the bottleneck (Lynch et al., 1995). This means that deleterious alleles may even increase in frequency in captivity and eventually fix (Lande, 1994; Whitlock, 2000), possibly increasing extinction risk (Lynch and Gabriel, 1990). However, the unmasking and subsequent expression of recessive deleterious alleles also exposes them to selection and, in theory, allows their removal in a process known as purging. Theory suggests that in most cases only highly deleterious alleles can be purged effectively during bottlenecks (Kirkpatrick and Jarne, 2000) but it is unknown to what extent purging can contribute to the success of captive breeding programs.

Purging is emerging as a possible contributor to the persistence of wild bottlenecked populations and has been detected in many vertebrates (Dussex et al., 2021; Foote et al., 2021; Grossen et al., 2020; Kleinman-Ruiz et al., 2022; Lan et al., 2023; Mathur et al., 2022; Robinson et al., 2018; von Seth et al., 2021; X. Wang et al., 2023; Xue et al., 2015). In general, large effect alleles tend to be purged more effectively whereas moderate and low impact alleles escape this process and may even fix (Xie et al., 2022), consistent with theoretical predictions (Kirkpatrick and Jarne, 2000). Purging has even been detected in the captively reared Dama gazelle (*Nanger dama*) which has an estimated effective population size of 11 (López-Cortegano et al., 2021). This is surprising, given that there appear to be limits to the ability of selection to purge alleles at extremely low effective population sizes (Glémin, 2003). While there is a growing body of literature examining the impact of purging on wild populations, few studies have turned to captively bred populations which often go through more extreme bottlenecks than those seen in the wild. An exception is the Chatham Island black robin, which experienced successive bottlenecks in the 20^th^ century, eventually being reduced to a single breeding pair in the wild. A pedigree study of the now recovering population found no evidence for purging of deleterious alleles (Kennedy et al., 2014). The ability of purging to act on deleterious alleles in these extreme cases is therefore still unclear. Where purging has been detected in captively bred populations (López-Cortegano et al., 2021) the analysis was borne out with pedigrees and phenotypic measurements. An investigation of allele frequency dynamics in such cases would enrich our understanding of the mechanistic basis of purging in captively bred populations.

In this study, we examine the molecular evidence for purging in the critically endangered *Dryococelus australis*, the Lord Howe Island stick insect. The species was once found abundantly on Lord Howe Island (LHI), 600 km east of Australia in the Tasman Sea, but was driven extinct by the accidental introduction of black rats (*Rattus rattus*) to LHI in 1918. In 2001, an expedition to the nearby sea-stack, Ball’s Pyramid (BP), discovered a remnant population (Priddel et al., 2003). In 2003 two mating pairs were removed from the island and used to successfully establish a breeding colony at the Melbourne Zoo (Carlile et al., 2009). Inbreeding depression is apparent in the captive population (Honan, 2008) and the BP population is small (Carlile et al., 2009). Previous investigators have suggested a genetic analysis of the species’ extant populations for these reasons (Carlile et al., 2009; Honan, 2008; Priddel et al., 2003). We sequenced the genome of one wild individual to investigate genetic diversity in the wild and found extremely low heterozygosity and long runs-of-homozygosity suggestive of recent inbreeding events. We then leverage low-coverage whole genome sequence data from captive and wild *D. australis* to investigate the effects of the captivity bottleneck on genetic diversity. By approximating the fitness effects of alleles by their predicted impact on protein function, we were also able to examine the evidence for purging.

Wild individuals have high absolute inbreeding which is only exacerbated by captivity. Only the most highly deleterious alleles, frameshift and stop-codon mutations, show a greater loss in captivity compared to the wild. All other allele types have similar frequency changes, regardless of their predicted fitness effect. This suggests that purging has acted, but only on the most detrimental alleles. The stronger the predicted impact of an allele, the more likely it was to be found outside of runs-of-homozygosity (ROH), suggesting inbreeding rather than drift as the mechanism of purging. Our results suggest that purging can act even in extreme cases and complement those of López-Cortegano et al. (2021) who suggest that allowing breeding contributions to be governed by natural selection and random mating may facilitate the persistence of captive breeding populations in the long term. However, all such decisions must be weighed against the expected magnitude of drift, which will inevitably increase the genetic load of a population by randomly increasing deleterious allele frequencies.

## Methods

### Whole-genome resequencing

We generated low-coverage whole-genome sequencing data from 24 adult *D. australis*. Two of these were one male and one female offspring of an adult female collected from BP in 2017. This new female was gravid on collection, and so these two offspring are considered wild individuals. Hereafter, we refer to this female as the new founding female, and her two offspring as the wild male and female respectively. The other 22 adults were captive-born individuals from the Melbourne Zoo from two captive lines, LHISI and LHIP. The LHISI line is descended from the original founders of the captive breeding program whereas LHIP is a hybrid line derived from LHISI individuals and the wild male. In total, we sampled 12 LHISI individuals and 10 LHIP individuals, of even sex ratio. We refer to these three subsets of the data hereafter as the captive, hybrid, and wild groups. We generated further whole-genome sequence data for the wild male, who was thus sequenced at low and high coverage. All adults were sampled after dying of natural causes or euthanised in keeping with the approved terms of Zoos Victoria research project ZV21013.

DNA was extracted using Monarch gDNA Purification kits (New England BioLabs). We quantified extracts with the Qubit broad range DNA assay (ThermoFisher) and measured purity with a DropSense (Perkin Elmer) using standard 260/280 and 260/230 ratios. We also assessed DNA integrity by running 1 uL of extract on a 1.5 % agarose gel at 110 v for 30 minutes. Low-coverage samples were submitted to the Ramaciotti Centre for Genomics (Sydney, Australia) for PCR-free library prep and sequencing on an Illumina NovaSeq 6000 S1 flow cell (150 bp paired-end reads). An additional library was prepared in-house for the wild male with the NextFlex Rapid DNA-seq 2.0 kit (PerkinElmer), using half the recommended reaction volume. This library was sequenced at the Biomolecular Resource Facility (Australian National University, Canberra, Australia) on a NovaSeq 6000 SP flow cell (150 bp paired-end reads). Sequence data was delivered demultiplexed with adapters trimmed. Reads were aligned to the reference genome (Stuart et al., 2023) with bwa mem v0.7.17 (Li, 2013) and PCR duplicates removed with the samtools v1.12 markdup subroutine (Li et al., 2009). Mapping statistics are presented in Supplementary Table 1. As different library preparation methods were used to generate the low-coverage and high-coverage reads for the wild male, we did not pool them for analysis.

### Analysis of High-Coverage Sequence Data

We estimated mean coverage across the wild male’s genome in 100 kbp bins using the high-depth sequencing reads with MosDepth v0.3.1 (Pedersen and Quinlan, 2018). Mean depth across the autosomal genome was 20.5X (Supplementary Table 1, Supplementary Figure 1).

Following this, we called polymorphisms in the autosomal genome using freebayes v1.3.6 (Garrison and Marth, 2012) allowing a minimum mapping quality of 30, a minimum base quality of 20, and requiring a site to covered by a minimum and maximum of 10 and 30 reads to be considered. We retained homozygous reference calls in the resultant vcf file in order to correctly calculate genome-wide heterozygosity later. We filtered genotype calls first by removing any site falling into known repeat regions (Stuart et al., 2023) and then removed all indels. Finally, we used vcflib v1.0.10 (Garrisson et al., 2022) to remove sites with allele balances outside of the range of 0.2 – 0.8 and sites whose depth exceeded the genotype call quality by 4 or more. By splitting the resultant variants into heterozygotes and homozygotes we counted the number of filtered variants of both kinds falling in 1 mb sliding windows with an overlap of 500 kbp using bedtools v2.30.0 (Quinlan and Hall, 2010) and used this to calculate normalised heterozygosity for each window. Many regions of the genome were filtered out by the repeat mask and so our filtered variants represent on average 15 % of the entire genome. To avoid artefacts of low sampling density, we excluded genomic windows where fewer than 10 % of sites were retained after filtering.

To investigate the effect of variable recombination rate across chromosomes, we calculated the mean heterozygosity of each autosomal scaffold using the values calculated for the non-overlapping bins, weighting the calculation by the number of sites included in each bin. We took scaffold length as a coarse proxy for relative recombination rate (the per base recombination rate being necessarily higher in smaller chromosomes) and plotted this against mean heterozygosity. We also estimated the GC content of each of the wild male’s chromosomes. We first took the consensus sequence of our reference genome and all the wild male’s homozygous alternate SNP calls. We then used bedtools to generate 1,000,000 randomly located 100 bp intervals across the autosomal scaffolds (minus repetitive regions) and calculated the GC content of each using seqtk v1.3-r106 (https://github.com/lh3/seqtk). We calculated the mean estimated GC content of each autosomal scaffold using these windows and plotted this against scaffold length. We used these GC and heterozygosity values to estimate linear models for each in R v4.3.3 (R Core Team, 2018) using scaffold length as the predictor.

To compare genetic diversity in *D. australis* to other species, we collected genome-wide heterozygosity values estimated using whole-genome sequencing from the literature (Supplementary Table 2) and compared these to the genome-wide mean for the wild male.

We then used bcftools v1.19-5-g2bbfdbf9 (Danecek et al., 2021) to call ROH. As our sample consists of just one individual, we used default allele frequency values of 0.4, 0.5, and 0.6, to estimate the sensitivity of ROH calls to this parameter. We plotted tallies of the number and length of detected ROH using ggplot2 (Wickham, 2016) implemented in R.

### Analysis of Low-Coverage Sequence Data

#### Identifying Polymorphic Sites and Accounting for Assembly Error

We estimated mean sequence coverage in 100 kbp bins across the genome using MosDepth v0.3.1 and calculated mean depth of coverage across the autosomes and the X chromosome. Five individuals had < 1X mean autosome depth of coverage so we excluded them from further analyses. For the remainder, mean autosome depth ranged from 2.7-6.0X for the captive-born individuals (which includes the captive and hybrid groups) and 1.8-2.1X for the wild individuals (Supplementary Table 1, Supplementary Figure 2), and so we analysed the data with angsd v0.937-81-g88fd84d (Korneliussen et al., 2014), a software suite designed to analyse low-coverage sequence data by the calculation of genotype likelihoods. For this analysis and all subsequent analyses using angsd, we used the following filter: ‘-uniqueOnly 1 -remove_bads 1 -only_proper_pairs 1 -trim 0 -C 50 -baq 1 -minMapQ 30 -minQ 30’. This removes all sequences that did not map uniquely and mappings with a sam format flag greater than 255 (not primary, failure, or duplicate reads), retains only sequences whose mate mapped to the same scaffold and that map with a quality greater than 30 after adjusting the mapping quality score for mismatches, ignores bases of quality lower than 30, and uses the base quality alignment score used by default in samtools. We also only included sites not covered by the assembly repeat mask. This mask covers 2.2 Gbp of the 3.4 Gbp reference genome assembly. We also only considered sites on the autosomes, leaving 1.1 Gbp of the genome assembly available to this analysis. Other filters are specific to different angsd sub-routines and we describe them further below as necessary.

We identified segregating sites within populations and fixed differences between populations. First, we used the -doMaf subroutine of angsd to obtain per-site estimates of minor allele frequency across the autosomes excluding regions masked by RepeatMasker (Smit et al., 2021) (see Stuart et al. (2023) for a description of the repeat identification process). We set the major allele as the reference. Excluding the individuals who sequenced poorly mentioned above, we estimated allele frequencies for all individuals in all three groups, requiring a range of 2-8X coverage for an individual to be included at a given site and requiring data from at least 9 individuals for a site to be considered. In this way, we capture all sites (passing filters) that are segregating or fixed among the groups. Then, we estimated allele frequencies for each population separately at the sites detected across all individuals. For the hybrid and captive groups, we required the same 2-8X depth of coverage from at least 4 individuals for a site to be included, and the wild populations 1-5X coverage from both individuals. Requiring coverage in both individuals is undoubtedly a conservative filter, but we think this is appropriate given our sample size of 4 chromosomes per site for this group.

Using the polymorphic sites identified in the allele frequency analysis, we used ngsParalog (Linderoth, 2018) to identify regions of problematic mapping not captured by our repeat mask. This software uses a likelihood approach to model sequencing errors and within-individual read depth variation to detect regions of putative paralogy that may be misassembled, leading to signals of inflated depth and excess heterozygosity across samples. We combined the results of this analysis with our repeat mask in all downstream analysis to filter out sites and regions in our assembly that may bias results, and we refer to this hereafter as the combined mask. Overall, 3.2 % (35.4 Mbp) of the non-repetitive autosomal genome was covered by regions of putative mis-mapping, leaving 264,493 of 328,673 polymorphic sites for downstream analyses. We recognise that our analysis of mis-mapping is of a coarse grain, as ngsParalog can only make use of polymorphic sites in the genome, but due to the sparse nature of our data and the highly repetitive nature of the genome sequence, we believe a conservative approach to inference is appropriate.

#### Population Structure

To estimate the population structure among our samples, we used PCangsd v0.98 (Meisner and Albrechtsen, 2018). PCangsd estimates covariance matrices from sequence alignment files and is designed for use with low-depth, heterogeneously structured data. We filtered the list of polymorphic sites across all samples identified previously with angsd, removing sites contained in the combined mask. Additionally, we filtered sites with a minor allele frequency of 1/2N, where N is the sample size. Since we lacked the sample size to estimate linkage disequilibrium, we obtained lists of sites at least 50 kbp away from each other using the vcftools ‘--thin’ option with a dummy vcf containing site information as input. Then, we used angsd to estimate genotype likelihoods at these sites as input for PCangsd, using the same read filters described above, requiring a read depth of 1-8X in a minimum of 9 samples for a site to be considered. We used these genotype likelihoods to estimate sample covariance matrices with PCangsd, applying several additional minor allele frequency cutoffs (0.05, 0.10, 0.15) to observe their effect on sample clustering. We used R to estimate the eigenvectors of the covariance matrix and plotted samples in eigen-space.

#### Genetic Diversity

Next, we estimated the single sample folded site-frequency spectrum across all sites on each autosomal scaffold, removing sites falling inside the combined mask. We then calculated individual genome-wide heterozygosity as the fraction of sites with the site-pattern ½. We first performed this analysis using all non-repetitive, non-mismapped sites (with coverage for a given individual) as input, but the estimated values were strongly negatively correlated with mean sample depth. We reran analyses twice for all individuals, once using only sites at 2-4X coverage and using only sites at 5-8X coverage and found a much weaker relationship between depth and heterozygosity (Supplementary Figure 3). In this way, we controlled for depth differences between individuals in heterozygosity estimation. Whole genome heterozygosity was then calculated as the average of the separate scaffold estimates weighted by the number of sites included in each estimate, which varied depending on mean sample depth and the depth class used to create the input mask. We compared heterozygosity among the captive and hybrid groups with a *t*-test with unequal variances. One sample, C01220, a female from the hybrid group, was a clear outlier, and so we calculated the *t*-statistic including and excluding this individual. We did not include the wild group in any comparison due to the low sample size (N = 2).

#### Inbreeding and Runs-of-Homozygosity

Next, we explored the effects of the bottleneck on homozygosity throughout the genome. We used the R package RZooRoH v0.3.1 (Bertrand et al., 2019) to estimate the location and length of runs-of-homozygosity (ROH) in the hybrid, captive, and wild individuals. ROH are formed by inbreeding events and then broken up over time by recombination. Long ROH are formed by recent inbreeding events, while short ROH represent more distant events. In this way, the distribution of the lengths of ROH carry demographic information at multiple time scales (Ceballos et al., 2018). Identifying genome-wide ROH has proven to be an efficient and accurate method for characterising individual inbreeding (Alemu et al., 2021; Caballero et al., 2021; Keller et al., 2011; Nietlisbach et al., 2019), where the traditional F statistic based on comparing sub-population allele frequencies is replaced by the fraction of the genome contained within ROH, F_ROH_ (McQuillan et al., 2008). RZooRoH employs a hidden Markov model (HMM) to evaluate the probability that a given position resides within or without ROH in a given individual. Model-based approaches can more robustly estimate relative (individual or population level) inbreeding than window-based allele-counting approaches with reduced representation or low-coverage WGS data (Bertrand et al., 2019; Druet and Gautier, 2017).

We estimated genotype likelihoods for all individuals, using the same read filters as above and excluding the combined mask. Additionally, we imposed a depth filter of 1-8X for an individual to be considered for a given site, and a minimum coverage of 9 samples for a site to be considered. We then modelled ROH per-individual but using the same model parameters for all individuals simultaneously. This is recommended by the developers (Bertrand et al., 2019) to facilitate comparison among individuals. The pertinent model parameters are *K*, the number of ROH classes to model, and *R_K_*, where the probability of an ROH of class *K* ending between two SNP positions *d* Morgans apart is *e^−Rkd^*. Without changing default settings, one Morgan corresponds to 100 Mbp of sequence. High rates correspond to short (old) ROH and low rates correspond to long (young) ROH. There is a rough correspondence between a class’s rate and the inbreeding loop connecting the two ancestors whose mating produced that class’s ROH but such measures do not substitute direct knowledge of the pedigree. The model estimates the probability that each input position for each individual resides in one of *K* classes, accounting for genotype uncertainty and sequencing error. We explored the parameter space of *R_K_* and *K*, estimating per-individual models for all individuals for pairwise combinations of *R_K_ = {2, 5, 10}* and *K = {6,7,8,9,10}*, where each combination of parameters generates a set containing stopping rates for each class corresponding to *R_K_* raised to the power of the i^th^ class defined by *K*, e.g. the parameter combination *R_K_ = 2* and *K = 6* generates a set of rates from 2^1^ to 2^5^, with the 6^th^ class reserved for sites not in ROH.

We began by including genotype likelihoods for all individuals at once. This facilitates comparison of individuals from different populations with respect to the same base population but may bias our estimation of F_ROH_. Since the estimation procedure relies on allele frequencies calculated directly from the data, we estimated models for the captive and hybrid groups separately, using only the respective population’s genotype likelihoods as input. We then compared F_ROH_ for each individual, as well as the per-individual models’ Bayesian information criterion (BIC), across model parameters and the base population used as input. We found that, for a given base population, model parameters had a mostly negligible effect on F_ROH_, estimates slightly increasing with the addition of more ROH classes (Supplementary Figure 4).

We plotted and compared the distribution of F_ROH_ in our three groups. We tested the mean difference between the captive and hybrid groups with a *t*-test with unequal variances. We also compared in a similar manner the total number and total summed length of ROH in length classes of 0 – 10 kbp, 10 – 100 kbp, 100 – 300 kbp, 300 – 500 kbp, 500 kbp – 1 Mbp, and > 1 Mbp. The individual from the hybrid group with outlying heterozygosity also had outlying F_ROH_, and so we calculated test-statistics including and excluding it.

#### Characterising Deleterious Mutation Load

We used the allele frequencies estimated for all polymorphic sites identified previously by the -doMaf angsd subroutine. This included polymorphisms with respect to the captive, hybrid, and wild individuals, so any allele fixed in captivity but absent in the wild, or vice versa, was included. We annotated all polymorphisms with snpEff v5.0 (Cingolani et al., 2012), using our reference genome assembly and coding sequence annotation as input, and then removed any sites that fell within the combined mask, as allele frequencies were estimated as a first step prior to the identification of misassembled regions with ngsParalog. SnpEff annotates alleles by the severity of their predicted impact on protein function, with alleles in intronic sites and sites > 5 kbp up or downstream of a protein coding gene being classed as ‘modifier’ alleles, which we refer to here as non-coding. Low impact alleles are synonymous polymorphisms. Moderate impact alleles are in-frame deletions (whole codon deletions) and missense non-synonymous polymorphisms. High impact alleles are frameshift and stop-codon alleles.

With this data we then estimated the relative deleterious allele load in the hybrid and captive groups. We counted the number of sites carrying segregating or fixed alleles of different effect classes in the hybrid, wild, and wild groups and compared the counts between populations with a χ^2^ test. We identified alleles detected in the wild individuals and the hybrid and captive groups and calculated the frequency difference for each of these for the respective groups. In the hybrid group, these alleles are directly descended from the wild male. In the captive group, these alleles are descended from the two original founders taken from the wild in 2003. We infer from this that these alleles are segregating at high frequency in the wild population, having been sampled independently on two separate occasions (in 2003 and 2017). We compared the frequency differences of the different effect classes (high, moderate, low, and non-coding) to estimate their persistence through the bottleneck with respect to the expected strength of selection based on the snpEff annotations. We then analysed these differences with simple linear models in a Bayesian framework using brms (Bürkner, 2017). Two sets of models were run, one where the allele effect classes were coded as an ordered factor (from non-coding to high impact) and one where effect class was unordered and so a slope parameter was estimated for each class separately. In both model types, the non-coding effect class was treated as the reference, and so slope terms represent the difference of the other allele classes with respect to the non-coding one. The captive and hybrid subpopulations were modelled separately to avoid pseudoreplication of observations at the locus level. For each model, we ran four chains of 3,000 steps and discarded the first 1,000 steps as burn-in. Once models were complete, we ran hypothesis tests estimating the posterior probability that the slope term (either for each allele class or the estimates for linear, quadratic, and cubic effects in the case of the ordered factor models) was less than 0 since we have the *a priori* expectation that the frequency change in captivity should be more negative for alleles in coding regions compared to coding regions. Then, to approximate the contribution of inbreeding to the removal of deleterious alleles from the populations, we extracted the coordinates of all ROH identified previously with RZooRoH, using the *K* = 9 and *R_K_* = 2 model including all individuals simultaneously, and classified each site by the proportion of individuals in each of the hybrid and captive groups in which the site was contained in ROH. We plotted the percentage of segregating alleles of different effect classes that were contained in ROH in < 50 % and ≥ 50 % of individuals and generated 95 % confidence intervals around these percentages by performing 100 bootstraps with resampling. For each effect class, a bootstrap sample consisted of a random sample with replacement of all sites in that effect class, sampling as many sites as in the original dataset.

The annotations used here are of varying quality, indicated by the low BUSCO score of the protein annotations (Stuart et al., 2023). It is possible that some of our results may be driven by low-quality annotations. To account for this, we reran the above analyses of allele frequencies while filtering out low annotation edit distance (AED) scores. AED scores measure the discordance between the annotations derived from transcript alignment and from protein homology and range from 0 to 1, 0 being the preferable score where there is no discordance between the two information sources. We reran our deleterious mutation analyses excluding sites falling into proteins with AED scores greater than 0.8, 0.7, and 0.5. Since our sample of segregating sites is small, it is possible that this filtering might also impact our results simply by reducing signal via decreased sample size. Therefore, we reran analyses again but excluding AED scores less than 0.2, 0.3, and 0.5. In this way, we were able to determine if low-quality annotations were driving the pattern of our results.

## Results

### Resequencing

Excluding the five poorly sequenced individuals, low-coverage sequencing from the captive and hybrid groups produced an average of 110.6 million reads (82.6 – 173.5), and of these an average of 88.6 % (85.9 – 90.1) were mapped to the reference genome following duplicate removal. The two wild individuals had lower sequencing yields overall, but not low enough to consider exclusion. Among the captive and hybrid samples, the mean per-base coverage across the autosomal scaffolds was 3.9X (per-individual means ranged from 2.73 – 6.03); the male and female wild samples had mean autosomal coverage values of 1.8X and 2.1X respectively (Supplementary Table 1, Supplementary Figure 2). The high coverage sequencing of the wild male produced 809,439,612 reads, 70.29 % of which were non-duplicate and mapped successfully, resulting in a mean coverage of 20.5X across the autosomes (Supplementary Table 1, Supplementary Figure 1). Mean depth for each autosome was similar but there was variation in the upper limits of the depth distribution, some chromosomes having much higher maximum depths than others (Supplementary Figure 1).

### Analysis of High-Coverage Sequence Data

The landscape of variation in the wild male was punctuated by large regions of exceptionally low heterozygosity with the largest of these being found on scaffolds CM056995.1 and CM056996.1 (Figure 1a). Whole genome heterozygosity was 0.036 % (Figure 1a). Mean heterozygosity varied by more than two-fold among scaffolds, with a minimum and maximum of 0.021 and 0.056 % respectively (Figure 1b). There was a significant negative relationship between scaffold heterozygosity and length (Figure 1b) (*F* = 5.069, df = 14, P = 0.041), heterozygosity decreasing in the longest scaffolds. This was also the case for GC content (Figure 1c) (*F* = 14.980, df = 14, P = 0.002). Compared to the published estimates of genome-wide heterozygosity from other metazoans, the wild male was the sixth lowest overall (Figure 1d), lower than any other invertebrate included in the dataset. The species with lower estimates were, in order from highest to lowest heteryzogisity, the Tasmanian devil (Miller et al., 2011), the snow leopard (Cho et al., 2013), the African cheetah (Dorbrynin et al., 2015), the Californian island foxes (Robinson et al., 2016; presented here using the mean of all individuals sequenced from the cited study), and the baiji (Zhou et al., 2013). These are all examples of species which are critically endangered or functionally extinct.

**Figure 1.**
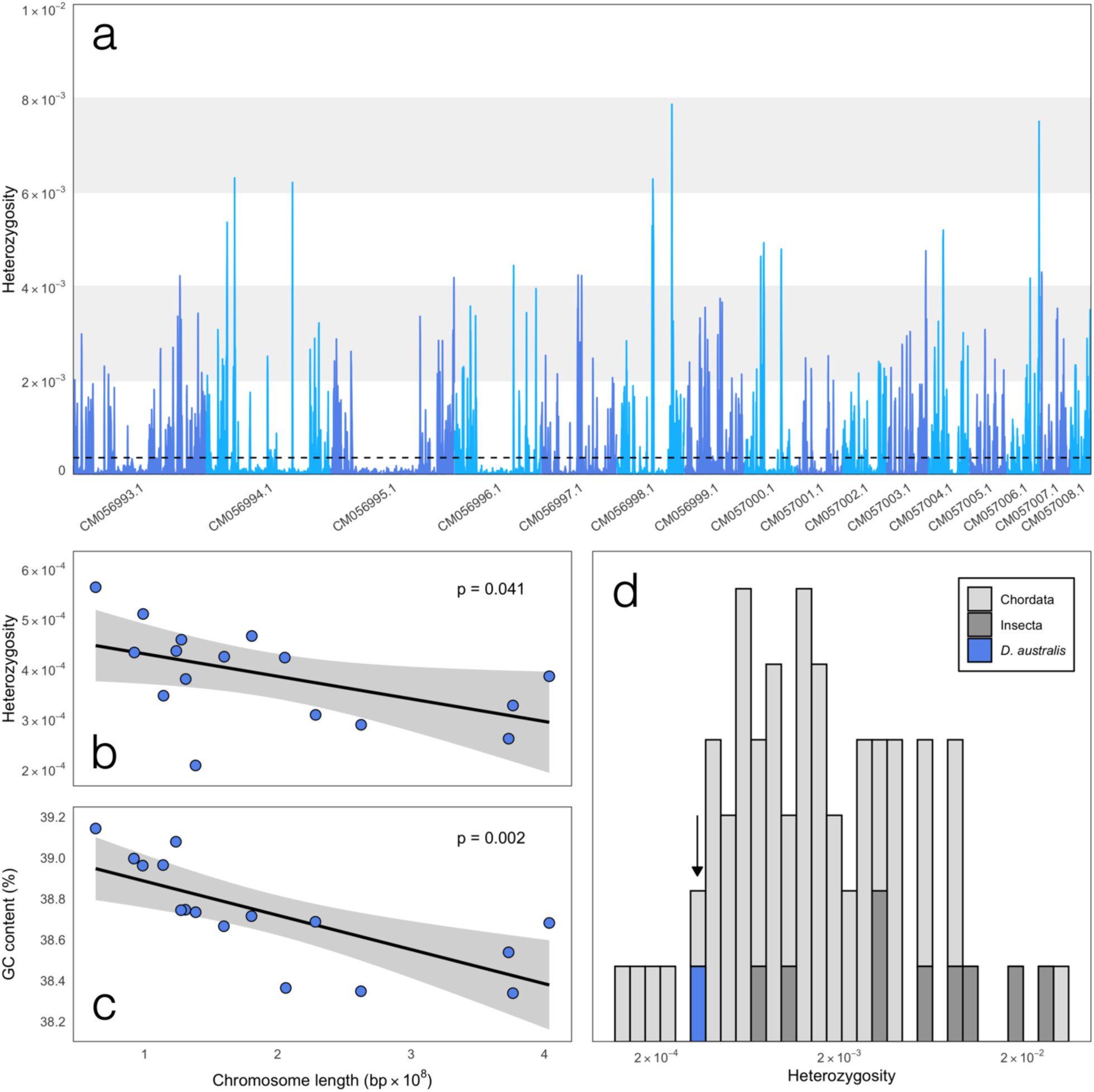
Genome-wide heterozygosity in a wild sample of *Dryococelus australis*. (a) Distribution of genome-wide heterozygosity in 1 Mb sliding windows with a 500 kb overlap across autosomal scaffolds. Polymorphic sites were called using freebayes freebayes v1.3.6 (Garrison and Marth, 2012). Dashed line represents the mean across non-overlapping windows (i.e. every second window), weighted by the total number of polymorphic and monomorphic sites included in that window after depth and quality filtering. (b,c) Relationships between autosomal scaffold length and heterozygosity (b) and GC content (c) respectively. Heterozygosity for each scaffold was calculated as for the genome-wide value in (a). GC content was estimated from 1,000,000 randomly generated 100 bp windows across the autosomes. Linear models were estimated in R v4.3.3 (R Core Team, 2018). Relationships for both heterozygosity (*F* = 5.069, df = 14, P = 0.041) and GC content (*F* = 14.980, df = 14, P = 0.002). (d) Comparison of genome-wide heterozygosity in *D. australis* with other published metazoan estimates. Bold arrow highlights the location of *D. australis*. Note that the x-axis is presented in the log scale.

The number of ROH called by bcftools increased with the allele frequency used and this was consistent across ROH length classes (Supplementary Figure 5). Qualitative results were identical for all three allele frequency classes, so we only report the results for an input allele frequency of 0.5, although we present some results for all three values in figures (Figure 2, Supplementary Figure 5). The greatest proportion of each scaffold in ROH was found on scaffold CM057001.1 followed by CM056996.1 and CM056995.1, however the longest ROH were found on CM056996.1 and CM056995.1, which were 8,031,431 bp and 7,931,340 bp respectively. This qualitatively agrees with the results of the sliding window analysis of heterozygosity although the total apparent length of the low heterozyosity regions (Figure 1a) is much larger than the maximum detected ROH lengths. It is possible that these already large ROH originate from a much larger ROH generated by recent inbreeding, although the distances between the longest and second longest ROH detected for each of these two scaffolds was 15,868,761 and 6,158,383 bp. Mean ROH length across scaffolds varied from 928,818 to 1,567,986, with a grand mean of 1,239,938 ± 171,998 (standard deviation). Like other results, the estimated inbreeding coefficient F_ROH_ also increased with the allele frequency input to bcftools. Pemberton et al. (2012) noted that short ROH could be reflective of background patterns of linkage disequilibrium throughout the genome rather than contemporary or ancestral inbreeding, and so we show the final calculated F_ROH_ using different minimum length cutoffs. Using cutoffs of 100 kb, 1 Mb, 2 Mb, and 5 Mb, the estimated value was 0.45, 0.35, 0.19, and 0.03 respectively (Figure 2).

**Figure 2.**
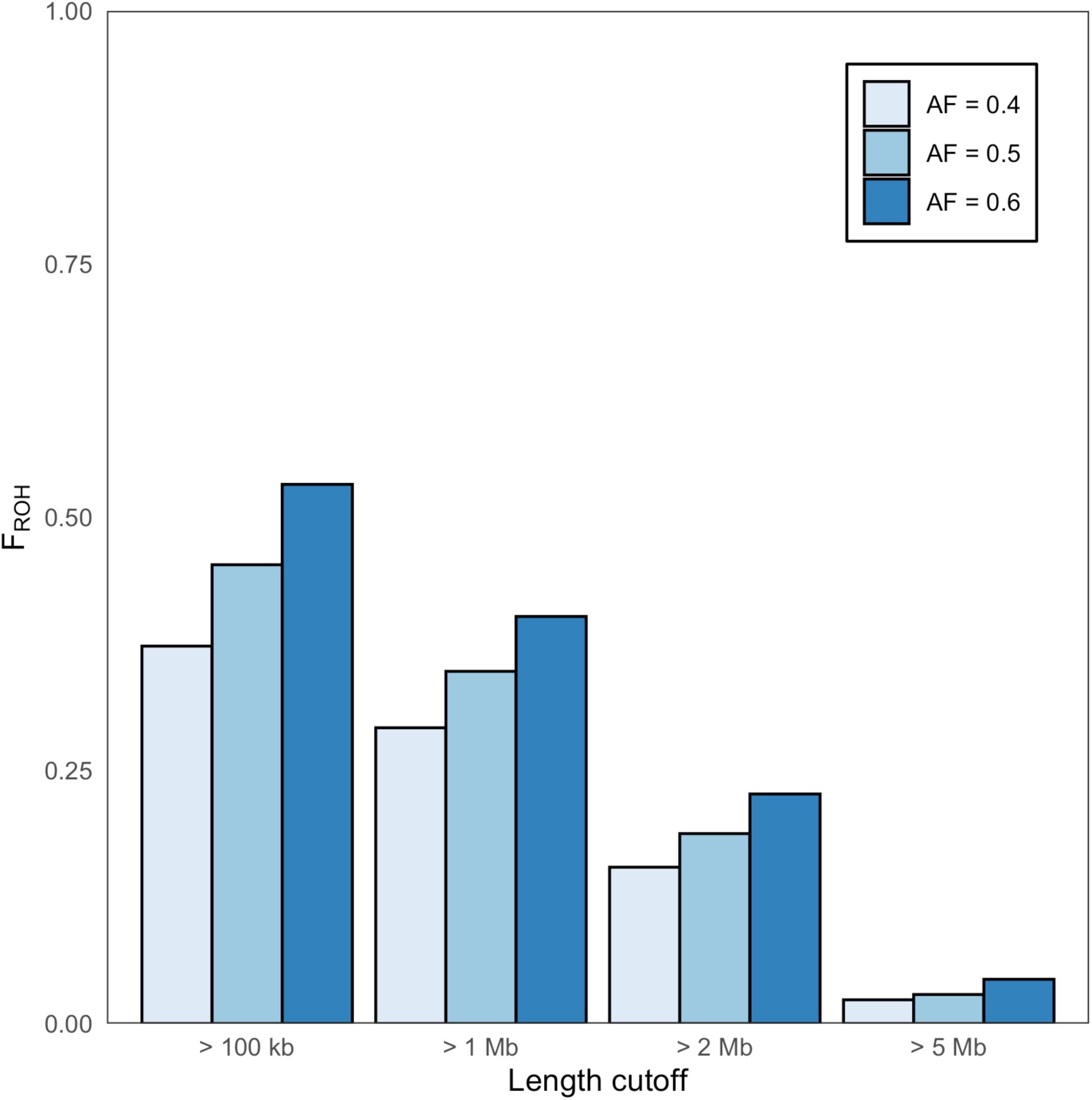
A large proportion of the wild *Dryococelus australis* genome is covered by runs-of-homozygosity (ROH) of varying lengths, suggesting both recent and historical inbreeding. ROH length and coordinates estimated using bcftools v1.19-5-g2bbfdbf9 (Danecek et al., 2021). Multiple starting allele frequency values were used for bcftools indicated by the colour legend. Different minimum length cutoffs are compared for each starting allele frequency.

### Analysis of Low-Coverage sequence Data

#### Population Structure

We used PCAngsd to estimate the covariance of allele frequencies in our low-coverage sample. Clustering across all 19 included samples was similar regardless of the minor allele frequency threshold applied (Supplementary Figure 6). The first two principal components of the covariance matrix estimated from all thinned sites with a minor allele frequency cut-off of 0.1 explained 26 % and 7 % of the covariance among samples. Wild individuals occupy one region of eigenspace defined by the first PC, the captive group another region, and the hybrid individuals all fall in between (Figure 3b), corresponding to the known population history (Figure 3a). Individual C01220, the low heterozygosity outlier, falls much closer to the captive individuals in the first PC than the other hybrid individuals.

**Figure 3.**
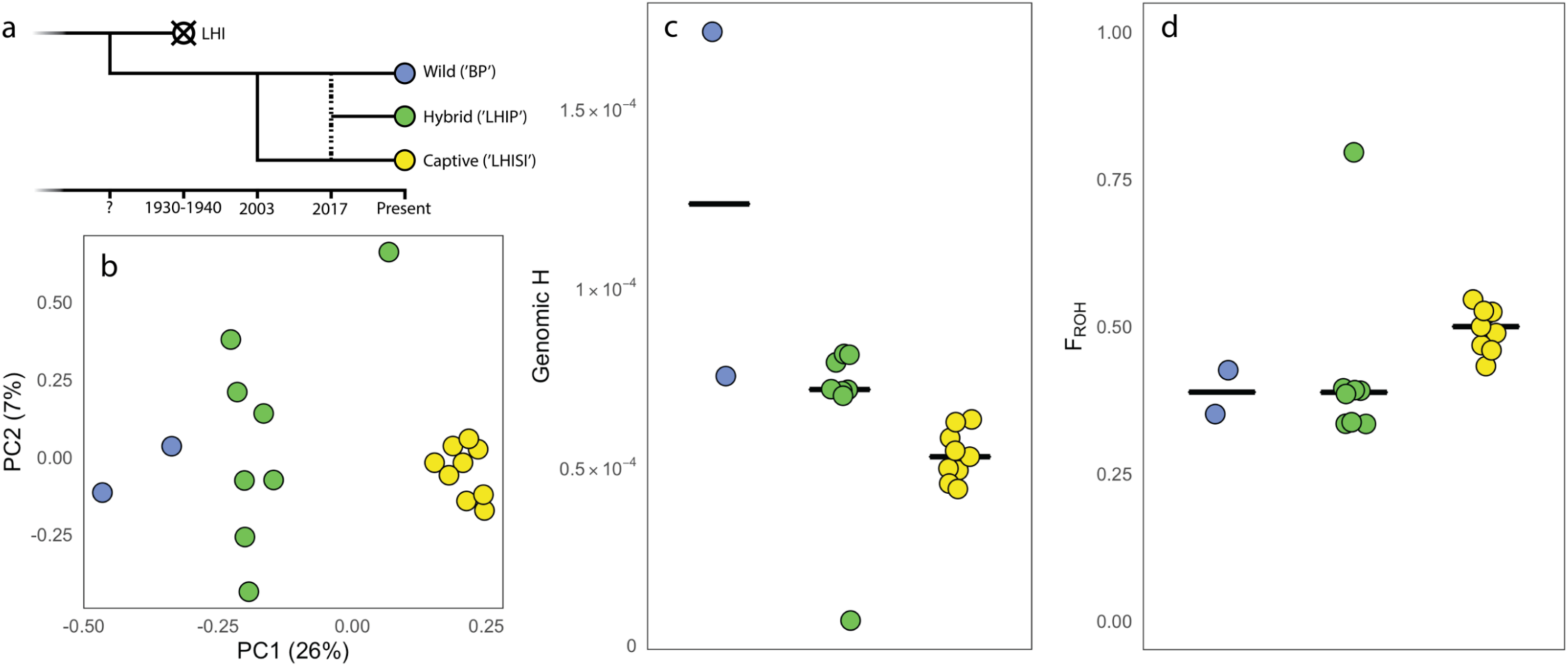
Recent demography has strongly depleted genetic diversity and increased inbreeding in *D. australis*. (a) Visual model of population demographic history of the sampled individuals. At an unknown time point (point ‘?’) the Ball’s Pyramid and Lord Howe Island populations diverged. Sometime between 1930-1940 the Lord Howe Island population went extinct after the introduction of rats to Lord Howe Island (Priddel et al., 2003). In 2003, the captive group was founded by taking a male and a female from the Ball’s Pyramid population. In 2017, another female was taken from Ball’s Pyramid, and her offspring were mated with captive individuals to produce the hybrid group. All three of these groups persist today. For (a), (b), and (c), the outlier in the hybrid group is individual C01220. (b) Population structure. Covariance matrix estimated using PCangsd v0.98 (Meisner and Albrechtsen, 2018) with a minor allele frequency cutoff of 10 % and eigenvectors calculated in R (R Core Team, 2018). (c) Autosomal heterozygosity calculated by estimating the single-sample site frequency spectrum for each sample using the angsd -doSaf subroutine (Korneliussen et al., 2014) across all sites at 5-8X depth of coverage. See Supplementary Table 3 for raw values and comparison of estimates at 5-8X and 2-4X depth of coverage. (d) Inbreeding coefficient calculated using the genome-wide probability of a position falling in a run of homozygosity estimated with the RZooRoH R package (Bertrand et al., 2019). The model presented here includes all samples for allele frequency estimation and *K* and *R_K_* values of 9 and 2.

#### Genetic Diversity

We calculated heterozygosity across all autosomal scaffolds for the 19 individuals with a mean coverage > 1X, using angsd to calculate the single-sample site frequency spectrum across all sites not excluded by our combined mask. This includes the two wild individuals, one of whom has a mean coverage lower than 2X, the minimum required to detect heterozygosity. However, even this individual has many individual sites with ≥ 2X coverage. We analysed low (2-4X) and moderate (5-8X) depth sites separately, due to the strong collinearity of mean sample coverage and estimated heterozygosity observed in preliminary analyses (not shown). To examine this artefact in our results, we plotted the number of sites analysed against estimated heterozygosity and found no significant relationship between the two for both the low and moderate depth data (Supplementary Figure 3). This suggests that normalising for depth differences across individuals is sufficient to remove this artefact. We excluded the two wild individuals from these figures as they are expected to be outliers in both mean sample depth and heterozygosity. Here, we present only figures from the moderate depth sites. See Supplementary Table 3 for full results. Individual C01220, a female from the hybrid group, had the lowest heterozygosity and the wild male had the highest (Figure 3c) at 0.017 %. This is considerably lower than the estimate of 0.036 % from high coverage sequencing (Figure 1a) suggesting that values derived from genotype likelihoods are probably underestimates. *Dryococelus australis* females may reproduce by parthenogenesis, and we consider this to be the most likely explanation for C01220’s low heterozygosity, which is further supported by her high inbreeding coefficient (see below, “Inbreeding and Runs-Of-Homozygosity”). Unless mentioned explicitly, this individual is excluded from all presentation of summary statistics in the text below, although we report the results of these tests both including and excluding her in the Supplementary Material (Supplementary Table 4a, 4b). The wild female had an estimated heterozygosity comparable to the hybrid group (Figure 3c). The captive group had clearly lower heterozygosity than the hybrid group (Figure 3c) (*t* = 7.171, df = 13.994, P < 0.001).

#### Inbreeding and Runs-of-Homozygosity

Modelling choices had variable impacts on the estimation of F_ROH_ and on the length and number of ROH, although overall trends across the hybrid and captive groups were consistent. Estimating ROH simultaneously for all individuals upwardly biased F_ROH_ in most individuals from both the hybrid and captive groups (Supplementary Figure 7) and this bias ranged from −0.07 – 0.03 and 0.02 – 0.1 in the two groups respectively. Overall, model log likelihoods were higher per individual for the group-specific models. However, to compare individuals, we focused on estimating ROH across all groups simultaneously. Comparing model BIC values revealed almost no difference in model fit where *R_K_ = 5* and *R_K_ = 10* and F_ROH_ also did not differ between these models (Supplementary Figure 4). BIC was maximised at *K = 9* for most individuals when *R_K_ = 2* and so we proceeded with these values.

F_ROH_ varied between the three groups in our sample. The median F_ROH_ was highest in the captive group (0.50), and the same in the hybrid (0.39) and wild (0.39) groups (Figure 3d). Similar to the heterozygosity estimates, the hybrid C01220 is an outlier here, with an estimated F_ROH_ of 0.80. The wild male and female had very different estimated F_ROH_ values, 0.35 and 0.43 respectively. When excluding the hybrid outlier in the calculation, mean F_ROH_ was significantly different between the hybrid and captive groups (*t* = −7.77, df = 13.97, P < 0.001). This result was the same when comparing F_ROH_ estimated using the group-specific models (*t* = −3.81, df = 12.29, P = 0.002) (Supplementary Figure 8). F_ROH_ in the wild male was similar when using low coverage and high coverage sequencing. The estimate from genotype likelihoods using RZooRoH was 0.35, the same as the value estimated using high coverage genotypes and excluding short ROH < 1 Mb.

Detected ROH frequently overlapped between individuals, such that in some regions, the haplotypes detected in the hybrid individuals represented a mix of the captive and wild haplotypes. For example, a 15 Mbp region in the center of CM056998.1 harbours a long ROH in all captive individual (the ends of which are detected at different locations in different individuals); this ROH is absent in the wild individuals but only some hybrids have it (Figure 4a), commensurate to their mixed ancestry. In many cases, a single ROH in one individual was detected as two shorter ROH in another (Figure 4a). With denser data and higher coverage, it is likely that many of these smaller ROHs would be merged into a single large contiguous ROH common to multiple individuals.

**Figure 4.**
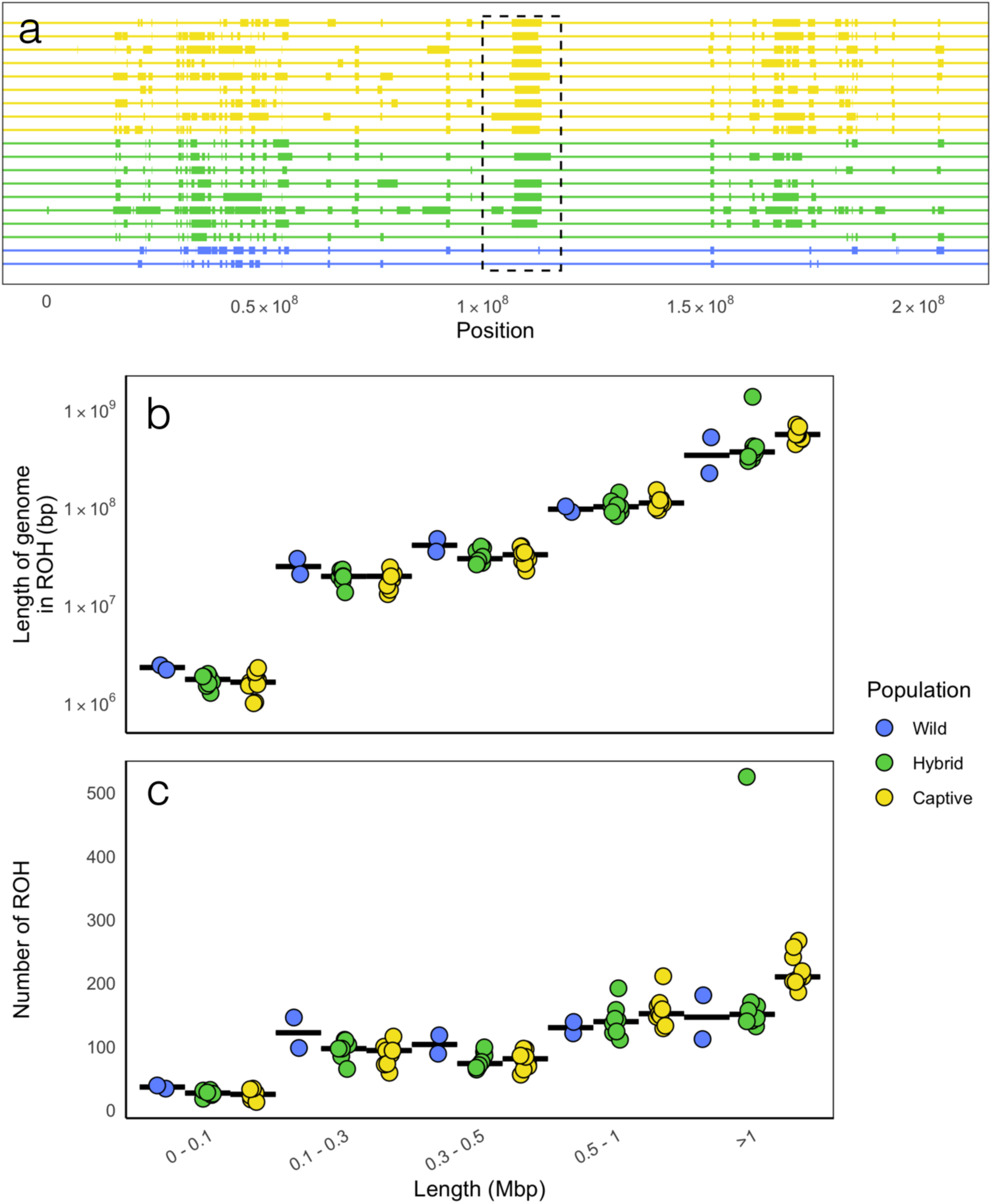
Runs-of-homozygosity estimated with the RZooRoH package (Bertrand et al., 2019) reflect recent and distant demography. The model presented here includes all samples for allele frequency estimation and K and R_K_ values of 9 and 2. (a) Predicted ROH on CM056998.1. Thin lines show the full length of the scaCold and thick lines demarcate the start and end positions of ROH. Black hashed box outlines a 15 Mbp region where hybrid individuals show a clear mixture of captive and wild haplotypes. (b) Cumulative length and (c) total number of ROH per individual across five length classes with black lines group medians. In (b) and (c), the outlier to the hybrid group in the ‘> 1 Mbp” length class is individual C01220.

Individuals in all three groups harboured many more long (> 500 kbp) than short ROH (Figure 4c) and the cumulative length covered by these long ROH was greater than for small ROH (Figure 4b). The captive group harboured more long ROH than the hybrid and wild groups, and there was a trend towards increased numbers and cumulative length of short (< 500 kbp) ROH in the wild individuals (Figure 4bc), although we did not test this difference due to the small sample size of this group. Again, the hybrid individual C01220 was an outlier with respect to both the number and cumulative length of long ROH (Figure 4bc). We tested the difference between the number of ROH of different size classes as well as the log cumulative length per individual between the hybrid and captive groups. Results mirror those of the tests comparing F_ROH_ and heterozygosity. The mean number of long ROH was greater in the captive than the hybrid group (500 kbp – 1 Mbp: *t* = 2.29, df = 13.70-, P = 0.04; > 1 Mbp: *t* = 6.76, df = 12.40, P < 0.001). The mean log cumulative length of long ROH was greater in the captive group (500 kbp – 1 Mbp: *t* = 2.47, df = 13.90, P = 0.03; > 1 Mbp: *t* = 6.41, df = 13.50, P < 0.001) (Supplementary Table 4a). There was no difference in means for neither number nor log cumulative length between the two groups for the other length classes (Supplementary Table 4a). This result was similar when considering the group specific models (Supplementary Figure 9, Supplementary Table 4b). In these cases, the exclusion of the hybrid outlier C01220 resulted in significantly different means for the greatest length class between the hybrid and captive groups, and regardless of the inclusion of C01220, the hybrid group had significantly greater mean numbers and log cumulative lengths of shorter ROH classes (Supplementary Table 4b).

#### Characterising Deleterious Mutation Load

In total, there were 264,493 polymorphic sites detected across all three groups, with the captive, hybrid, and wild groups carrying 188,020, 210,329, and 69,037 sites with detectable group-specific polymorphism (Supplementary Table 5). No group contained a significant excess of segregating alleles predicted to impact protein function compared to the others (χ^2^ = 5.452, df = 6, P = 0.487), although the direction of the residuals is potentially informative. The captive and hybrid groups carried an excess of impactful segregating alleles and the wild group carried a deficit compared to expectation, although we acknowledge that the wild sample size is very small. Across all groups, more alleles were annotated as moderately impactful than of low impact on protein function. For example, the captive group had 2,777 moderate and 1,276 low impact alleles segregating, a more than two-fold difference (Supplementary Table 5).

#### Detection of Purging

We analysed the frequency of non-reference alleles found in the two wild individuals compared to their frequency in the captive and hybrid groups to estimate the effect of purging on the persistence of alleles through the captivity bottleneck. For the hybrid group, the transmission of these alleles is direct. This is not the case for the captive group, therefore we excluded any alleles found in the wild but not the captive group in all analyses of the captive group, as we cannot distinguish between the case where alleles at these sites were transmitted via the original founders and lost over time and the case where such alleles were not present in the original founders. On the contrary, presence in the two wild individuals and absence in the hybrid group implies loss in transmission to the hybrid individuals, and so alleles found in the wild but not the hybrid group were retained for analysis of the hybrid group.

In both the hybrid and captive groups, the mean frequency difference of alleles shared with the wild individuals is more negative for the high impact class than the other mutation types (Figure 5a). We used linear modelling in a Bayesian framework to quantitatively assess the evidence for differences in frequency change among the classes. All chains converged and all *R̂* values reached 1 after 3,000 steps, and bulk and tail effective sample sizes were high (Supplementary Table 6a, 6b, 7a, 7b). For the ordered factor models, evidence for a negative effect of an allele’s effect class on frequency difference was strongest for the linear effects with successively weaker evidence for quartic and cubic effects (Supplementary Table 6a, 6b). Overall, the strength of evidence for negative linear effects of allele effect class on frequency change was stronger for the hybrid than the captive groups (posterior probabilities of 0.96 versus 0.91 for a slope < 0; Supplementary Table 6a, 6b). However, even for the hybrid group, the upper limit of the 95 % confidence interval for the linear slope term overlapped 0 (Supplementary Table 6b). Results for the non-ordered factor models were similar in that the strength of evidence for the effects of different allele effect classes was stronger for the hybrid (Supplementary Table 7b) than the captive group (Supplementary Table 7a). Taking 95 % posterior probability as a threshold for sufficient evidence for an effect, only the high effect alleles in the hybrid population showed evidence for a negative effect on allele frequency change in captivity, while the low and moderate classes showed similar strength of evidence (posterior probability = 0.69, 0.66; Supplementary Table 7b). For the captive group, evidence for a negative effect of effect class on frequency change increased from the low (posterior probability = 0.35) to the moderate (0.71) to the high (0.90) effect classes, consistent with the visual differences in mean frequency change (Figure 5a). In all cases, the modelled variance in allele frequency change (σ) was much larger than the mean change attributable to any effect class (Supplementary Table 6a, 6b, 7a, 7b) and this is also reflected in the visual distribution of frequency changes (Figure 5a).

**Figure 5.**
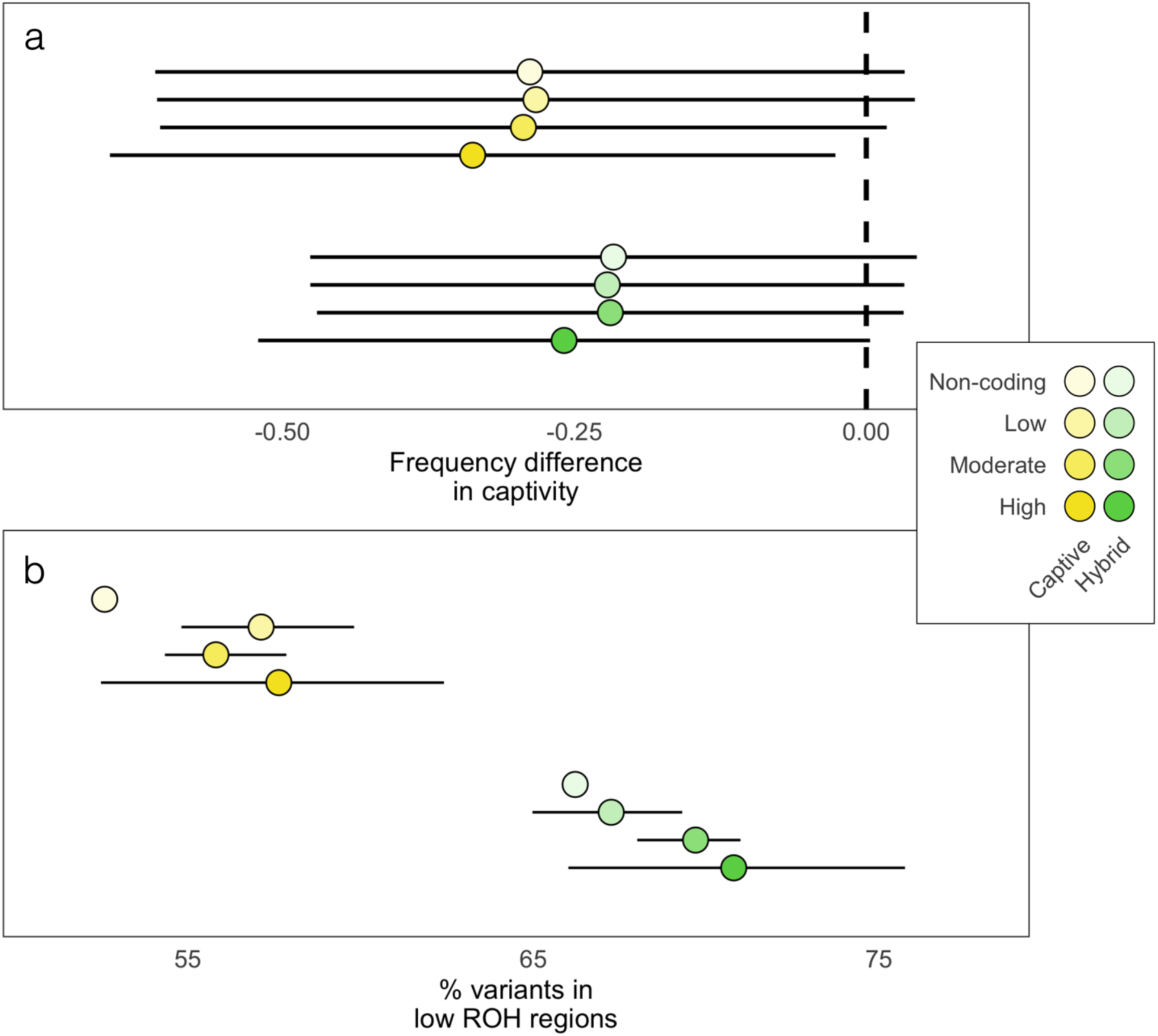
Highly deleterious mutations have been preferentially purged in captivity by inbreeding. Allele frequencies were estimated per group with the -doMaf subroutine in angsd (Korneliussen et al., 2014) and allele effects were estimated with snpEff (Cingolani et al., 2012). (a) Difference in frequencies between the two captive groups and the two wild individuals for segregating alleles. Points show mean difference and error bars show one standard deviation. Dashed line shows location of no change in allele frequency. The high impact annotation class has a more negative mean difference than the other classes. (b) Percentage of polymorphic sites in regions of low (< 50 %) ROH coverage with error bars showing 95 % confidence intervals of 1000 bootstraps. Non-coding alleles are more likely to be found in high (>= 50 %) ROH coverage than alleles with predicted impacts on protein function.

Next, we classified all alleles as falling into regions of high (>= 50 %) or low (< 50 %) coverage of ROH for both groups, and then calculated the percentage of segregating mutations falling into the two regions with bootstrapped estimates of 95 % confidence intervals. The number of segregating alleles of each class falling in regions of < 50 % ROH coverage generally increased from the non-coding to the high impact class for both groups, with more mutations of all classes falling into low ROH regions in the hybrid group due to the lower F_ROH_ (Figure 5b). In the captive group, none of the 95 % bootstrap confidence intervals for the low, moderate, and high impact mutation classes overlapped the mean percentage of non-coding mutations in low ROH regions (Figure 5b) The confidence intervals of the low and high impact classes overlapped the non-coding class mean percentage in the hybrid group. The breadth of the confidence intervals increased substantially from the non-coding to the high impact class due to sample size; the dataset comprises > 150,000 mutations in the non-coding class in both groups, but only a few hundred segregating high impact mutations were detected.

Our re-analysis which included AED score filtering showed no qualitative differences in the figures produced. Although there were subtle shifts in the means of allele frequency change for different effect classes, directionality of the differences between the effect classes was preserved (data not shown). This was the case when filtering out both low- and high-quality annotations.

## Discussion

### Purging Through an Extreme Bottleneck

We analysed the distribution of genetic diversity in wild and captively bred *D. australis* and found allele frequency patterns suggestive of purging through the captivity bottleneck. The frequency difference of highly deleterious alleles (frameshift and stop-codon mutations) was more negative than that of other mutation classes when comparing captive individuals and our small sample of wild individuals. When segregating in captivity, these highly deleterious alleles were more likely to be found in regions where fewer than 50 % of individuals harboured ROH, compared to non-coding segregating alleles of which roughly 50 % were in such regions. This implies the role of inbreeding in exposing recessive deleterious alleles to selection. This is despite a high rate of drift, implied by a substantial loss of heterozygosity in captivity. Heterozygosity was highest in a wild individual and lowest in individuals reared in captivity for 16 generations, but all individuals had low heterozygosity on an absolute scale.

These findings are consistent with early work on purging suggesting that only highly deleterious alleles are removed via purging (Hedrick, 1994; Wang, 2000; Wang et al., 1999). This has also been observed in wild populations such as alpine ibex (Grossen et al., 2020) which went through a severe domestication bottleneck, allowing the purging of highly deleterious alleles but the accumulation of moderately deleterious alleles. The post-bottleneck population increase results in an increased number of deleterious variants entering a population due to the increased opportunity for new mutation while simultaneously removing older, higher frequency deleterious variants (Gazave et al., 2013). In this way, the genetic load has a different distribution of fitness effects in stable compared to recently expanded populations (Lohmueller, 2014) even if the fitness burden of this load may be similar in the two (Simons et al., 2014). A more thorough sample of the wild BP population would allow us to test this finding in a wild species

Our comparison of alleles found in wild individuals and in the captive-bred individuals is indicative of purging against deleterious recessive alleles, although we acknowledge that the frequency of alleles across two closely related wild individuals is a poor estimate of the true wild allele frequency. However, by restricting our analysis to alleles segregating in these wild individuals and the hybrid and captive groups respectively, we restrict our analysis to alleles which must necessarily be common in the wild on BP, having been sampled in the former two individuals as well as in Adam and Eve, the population’s original founders. We observed a greater frequency difference between the wild individuals and the captive group for the most strongly impactful alleles (Figure 5a), but the posterior probability for an effect was only 0.90. This same difference resulted in a posterior probability of 0.96 for the hybrid group (Figure 5a). Any alleles present in the wild but not the captive individuals may not have been sampled in Adam and Eve, so they cannot be included in this analysis. This is not the case for the hybrid group. This creates a strong imbalance in sample size, restricting the statistical power for captive individuals. We also face the inherent limitation of a low diversity species. Coding mutations must necessarily be the least common mutations and in a highly inbred population these mutations would be even less prevalent. This means unavoidably low statistical power to detect mean frequency shifts between non-coding and coding mutations. We also observed a depletion of segregating coding alleles in regions where most individuals harboured ROH (Figure 5b); a greater proportion of segregating coding alleles were found in these low ROH regions than were segregating non-coding alleles. This last observation is independent of the estimation of allele frequencies in the wild individuals, and so is robust to the low wild sample size. We believe that the two observations, (1) of a depletion of coding alleles in captivity and (2) of their being preferentially located in regions with few ROH, strengthen each other. It is possible that selective sweeps may also contribute to the distribution of alleles within and without of ROH that we see here. Any beneficial allele that has experienced a selective sleep would be more likely to be found within ROH. It is difficult to speculate on exactly which effect class would be the most likely to harbour beneficial alleles, but it is probably the low and moderate impact classes, corresponding to synonymous, non-synonymous, and within-frame deletions. These changes are more likely to result in slightly altered protein function rather than non-function, as in the case of non-sense and frameshift mutations. This is another limitation of our method here, which assumes that any change to protein function has a deleterious fitness effect.

In the wild, it is likely that drift is an important process shaping genome wide diversity as it does for most of the mutation types analysed in this study. There may, however, be some influence of linked selection, whereby selection for or against non-neutral mutations reduces neutral variation in nearby regions of the genome which are physically linked (Kim and Stephan, 2000). Recombination allows neutral variation to become unhitched from these variants, and recombination rate variation can indeed explain a large proportion of variation in genetic diversity within a genome at a fine scale (Rettelbach et al., 2019). We found a negative relationship between scaffold length and heterozygosity (Figure 1b), consistent with the expectation under linked selection given that per base relative recombination rates are generally higher in shorter chromosomes (Kong et al., 2002). However, scaffold length was even more strongly positively correlated with GC content (Figure 1c), suggestive of GC-biased gene conversion, whereby recombination events favour the production of GC rather than AT gametes at polymorphic sites in the vicinity of double stranded breaks (Duret and Galtier, 2009). A negative correlation between chromosome arm length and GC content has been observed in humans as well (Katzman et al., 2011). Since both linked selection and GC-biased gene conversion are modulated by recombination rates, they can produce similar patterns of nucleotide diversity (Bolívar et al., 2015) which can be challenging to distinguish. It is not the express focus of this study to test competing hypotheses for the maintenance of genetic diversity in wild *D. australis*, but future work should follow up on this possibility if and when a larger sample of wild individuals is available.

One captive-bred individual appears to be the product of parthenogenesis rather than sexual reproduction. Although we cannot exclude unknown technical causes for low heterozygosity, C01220 was not an outlier in any of the quality control metrics that we examined, including mean sequencing depth, the number of sites included in the calculation of heterozygosity, read quality, read length, and the proportion of PCR duplicates among the raw reads. Based on the closely related *Extatosoma tiaratum* (Buckley et al., 2009, 2010; Forni et al., 2021), the most likely mode of parthenogenesis in *D. australis* is automixis with terminal fusion (Alavi et al., 2018). In cases of terminal fusion, heterozygosity is retained only at the distal ends of chromosomes, resulting in the loss of most of the mother’s genetic diversity. Individual C01220 has roughly 10 % of the heterozygosity of the other members of the hybrid group, consistent with this mechanism. This mode of parthenogenesis could enhance purging. Since most of the genome in an individual produced by automixis with terminal fusion is identical by descent, it can be considered an extreme form of inbreeding, even more so than selfing. This would accelerate the exposure of recessive alleles and facilitate purging and may have contributed to the effect detected here. This is an area where the development of theory describing the dynamics of allele frequency change in a population of mixed reproductive mode could be fruitful and lead to further interesting hypotheses to test.

### High Inbreeding in the Wild and in Captivity

We observed high F_ROH_ in all individuals sampled, although the median was higher for the captive group (0.50) than for the hybrid (0.35) and wild (0.36) groups. Is this level of inbreeding higher or lower than expected based on our knowledge of the captive population’s history? We can calculate the expected value of F, the traditional metric, in the captive group by making some simple assumptions under a coalescent framework. The average inbreeding in a population at time *t*, accounting for the level of inbreeding in the previous generation, can be calculated:

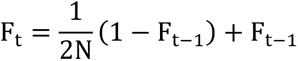

This simple coalescent model assumes no selection, constant population size, no additional mutation, and only sexual reproduction. If we take the median of the two wild individuals’ F_ROH_ estimates (0.36) and apply this model, we obtain an estimated F of 0.41, 0.46, and 0.54 for constant population sizes of 100, 50, and 25 respectively in the captive group after 16 generations. This is close to the median F_ROH_ in the captive group suggesting a very low effective population size in captivity over time. The wild *D. australis* F_ROH_ mirror those of other bottlenecked island populations, such as the kākāpō (Dussex et al., 2021) and the Sumatran rhinoceros (von Seth et al., 2021). Clearly, the history of *D. australis* is marked by recent and historical inbreeding events. The founding of the wild BP population may represent another bottleneck in the species’ history. Ball’s Pyramid and LHI are a short distance apart (23 km) but are separated by a large ocean trench (Standard, 1967) suggesting that the two islands may not have been recently adjoined. This would imply a very strong bottleneck in the population’s history as the colonisation of BP by rafting individuals is unlikely to have occurred multiple times. Alternatively, biogeographic analyses of the *D. australis* lineage suggest that it originated in Australia and migrated south through now submerged seamounts (Buckley et al., 2010). This would suggest that the BP and LHI populations were once continuous and were separated by gradual sea level change. This would create slow decline in population size rather than distinct bottleneck events.

We used RZooRoH (Bertrand et al., 2019) to detect ROH in our sampled individuals although we acknowledge that the use of reduced representation and lcWGS for ROH detection is still a fraught process (Duntsch et al., 2021; Lavanchy and Goudet, 2022). Despite differences in absolute estimates of F_ROH_ there is strong correlation between methods and data types (Duntsch et al., 2021) suggesting that relative comparisons are effective even with sparse data. Across samples, we expect there to be little bias introduced by differences in mean depth of coverage, as the wild individuals have the lowest mean depth (Supplementary Table 1) but the greatest number of short ROH (Figure 2bc). With incomplete representation of the genome, ROH detection methods tend to underestimate the number of short ROH (Lavanchy and Goudet, 2022), therefore the dropout of sites due to low depth in the wild individuals has not substantially biased the estimation of relative F_ROH_ and ROH tract lengths. The estimation of F_ROH_ with such approaches is less sensitive to data quality than is the determination of the precise coordinates of ROH (Duntsch et al., 2021; Lavanchy and Goudet, 2022). Apart from the wild male, all samples included in our analysis had a mean per-base coverage greater than 2, and so we expect most of the genome to have been sampled, although minimally for many sites.

We classified ROH into length classes. Going one step further and estimating age based on length is confounded by the inherent stochasticity of the meiotic recombination process by which large ROH become small over generational time (Thompson, 2013). Although such estimation has been performed for killer whale genomes (Foote et al., 2021) this analysis was based on high coverage genotype calls. We limit ourselves to a brief qualitative comparison. We observe large numbers of long ROH in all populations, and long ROH were the most prevalent in the LHISI group (Figure 2c). This is consistent with the effect of the captivity bottleneck on consanguinity. The presence of long ROH in all individuals can be attributed to recent inbreeding events in the wild on BP. The age of the wild BP population is unknown, but long ROH are consistent with a recent colonisation event from LHI. Mitochondrial divergence between BP and historical LHI specimens is low, and of a similar magnitude to divergence between different LHI specimens (Mikheyev et al. 2017), suggesting recent colonisation. All three groups had similar numbers and lengths of intermediate length ROH. These results are consistent with a long-term low population size. Another invertebrate from the region, the Lord Howe Island cockroach, *Panesthia lata*, also has populations on both LHI and BP. The divergence times between these have been estimated to at least 200 kya (Maxim Adams, unpublished manuscript) spanning multiple glacial cycles, implying a colonisation of BP when it shared a land bridge with LHI. In *D. australis* it is more likely that, given the abundance of long ROH in all individuals and the low mitochondrial divergence between LHI and BP, that the wild BP population has been roughly the same size since its recent founding, although this will require corroboration by more detailed genomic study of historical LHI specimens with which the founding of the BP population can be more accurately dated. Censuses of the wild BP population have never counted more than 40 individuals (Carlile et al., 2009) although the Pyramid’s topography makes thorough searching difficult.

### Extreme Loss of Heterozygosity

Autosomal heterozygosity derived from low coverage sequencing was exceptionally low in captivity, reaching a nadir of 0.004 % (excluding one outlier) (Supplementary Table 3), although we stress that this is likely an underestimate. Estimated genome-wide heterozygosity differed by a factor of two when comparing low and high coverage sequencing estimates in the wild male. However, even rescaling the low coverage values using the difference between the wild male’s estimates still leaves all individuals with low heterozygosity on an absolute scale. Heterozygosity was higher in the two wild individuals who respectively had autosomal heterozygosity of 0.0075 % and 0.017 % (Supplementary Table 3). These values are still well below reported genetic diversity across the metazoan tree of life (Supplementary Table 2). The heterozygosity values reported here are more reflective of studies of other small island populations such as the Californian channel island foxes (Robinson et al., 2016). This is supported by the comparison of the heterozygosity of the wild male derived from high coverage sequencing when compared with other published estimates (Figure 1d). Overall, the evidence supports extremely low wild population size, consistent with previous surveys (Carlile et al., 2009). Further study of the wild BP population may unveil greater heterozygosity than the pattern described here, which comes from few observations, and the two wild individuals may be half-siblings. The possibility of half-siblings comes from the almost two-fold difference in heterozygosity between the wild individuals (Figure 3c). *Dryococelus australis* is known to multiply mate in captivity (personal observations) and *T. cristinae* is known to be polyandrous (Arbuthnott et al., 2015) and so it is likely that while the two wild individuals share a mother, their fathers may be different.

## Conclusions

The deliberate application of purging in captivity is considered a risky strategy for the management of captive populations (Boakes et al., 2007; Leberg and Firmin, 2008). The small size of the wild BP population may have contributed to the successful establishment of the captive breeding program. Deleterious recessives accumulate in large populations where there is substantial opportunity for heterozygosis (Mathur and DeWoody, 2021). Smaller populations tend to harbour less of this “masked load”, which is converted into “realised load” when homozygosity increases (Bertorelle et al., 2022). Still, the first generations of the captive breeding program were subject to inbreeding depression, the eggs of F1 captives being considerably smaller than eggs lain by Eve, the foundress (Honan, 2008). There must therefore be at least some segregating deleterious recessive alleles on BP. We can speculate that while the most deleterious of these were removed during the bottleneck (see above discussion), the captive *D. australis* populations should now be accumulating moderately deleterious alleles which are likely to become problematic when reintroduction is attempted unless further founders are introduced from the BP population. Monitoring this diversity is challenging, and assays of neutral genetic diversity may not suffice to describe the effects of drift on functional variation (Teixera and Huber, 2021). However, patterns of neutral genetic diversity do still provide useful signals of demographic change, which themselves can act as indicators for the action of drift (Kardos et al., 2021). A genetic monitoring approach that makes use of neutral markers, or a mixture of neutral markers and markers closely linked to functional variation, would prove useful in the future study of *D. australis* captive population, allowing ancestry and demography to be tracked directly in this challenging species.

## Acknowledgements

We extend our thanks to the full invertebrate husbandry team at the Melbourne Zoo in Melbourne, Australia, for their past and ongoing work with the *D. australis* breeding program. O.P.S. was supported by an Australian Government Research Training Program scholarship. The low coverage sequence data was generated with support from the PhD student grant scheme of the Genetics Society of Australasia, funded by Illumina, Inc. Additional laboratory work was supported by the Ecological Society of Australia’s Holsworth Wildlife Research Endowment student grant, Australian Research Council Linkage Project LP210200654, and additional funding provided by Zoos Victoria. Sequencing and library preparation of the low coverage samples was performed by the Ramaciotti Centre for Genomics at the University of New South Wales, Sydney. Sequencing of the high coverage sample was performed at the Biomolecular Resource Facility, Australian National University, Canberra. All other laboratory work was performed at the Ecogenomics and Bioinformatics Lab at the Research School of Biology, Australian National University, Canberra

## Data Availability

All newly generated sequence data will be publicly available on the National Center for Biotechnology Information’s Sequence Read Archive under BioProject accession PRJNA1124435.

