## Supplementary Figures for "Purging of highly deleterious alleles through an extreme bottleneck"

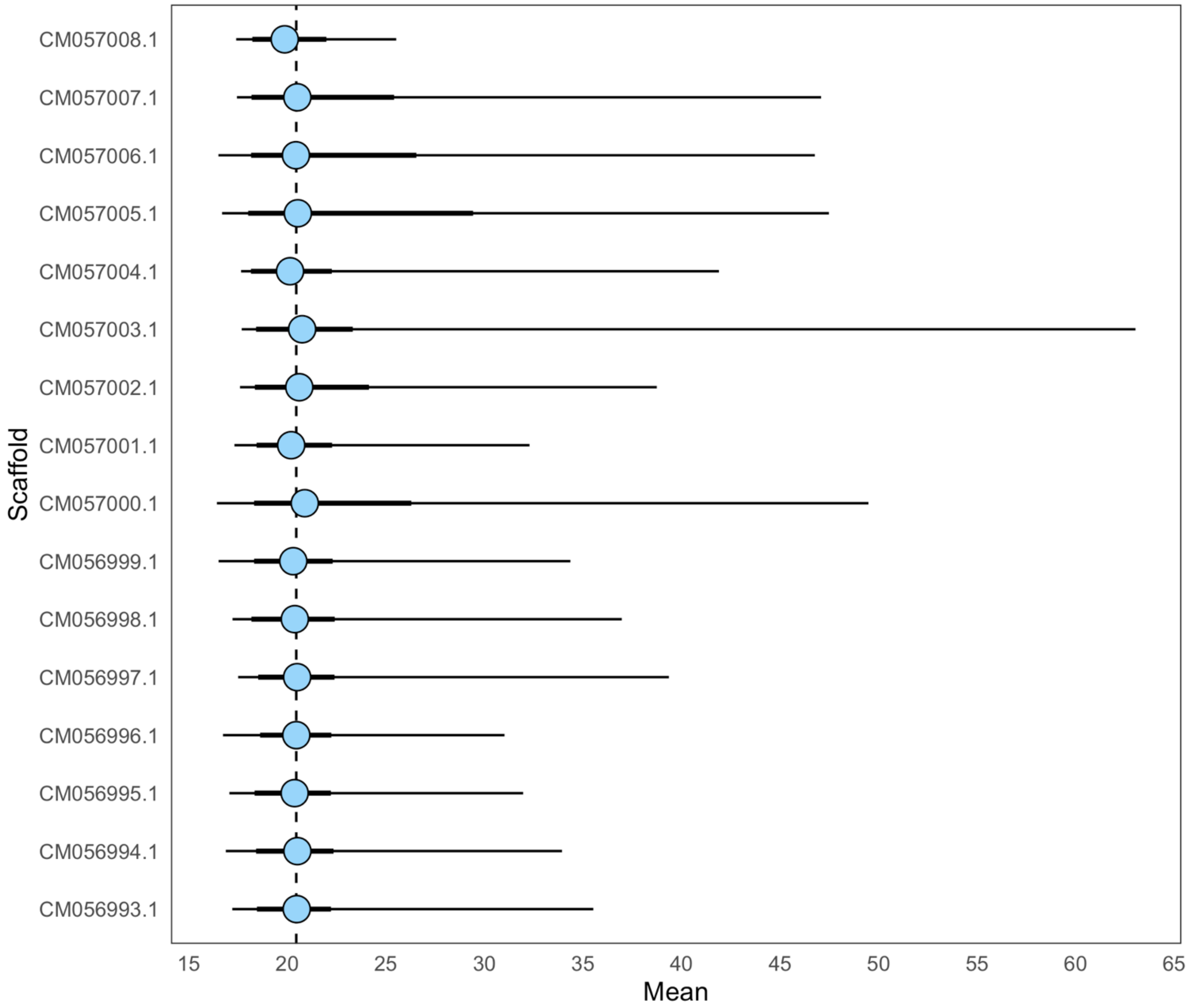


**Supplementary Figure 1.** Sequencing depth of coverage for the wild male's sequencing reads across the 16 autosomal scaffolds of the reference genome. Depth was calculated in 100 kbp non-overlapping bins using MosDepth v0.3.1 (Pedersen and Quinlan, 2018) For each scaffold, a mean is presented as well as the 95$^{th}$ (thick line) and 99$^{th}$ (thin line) percentiles of the coverage distribution. The dotted line represents the grand mean across all scaffolds, weighted by the length each scaffold.

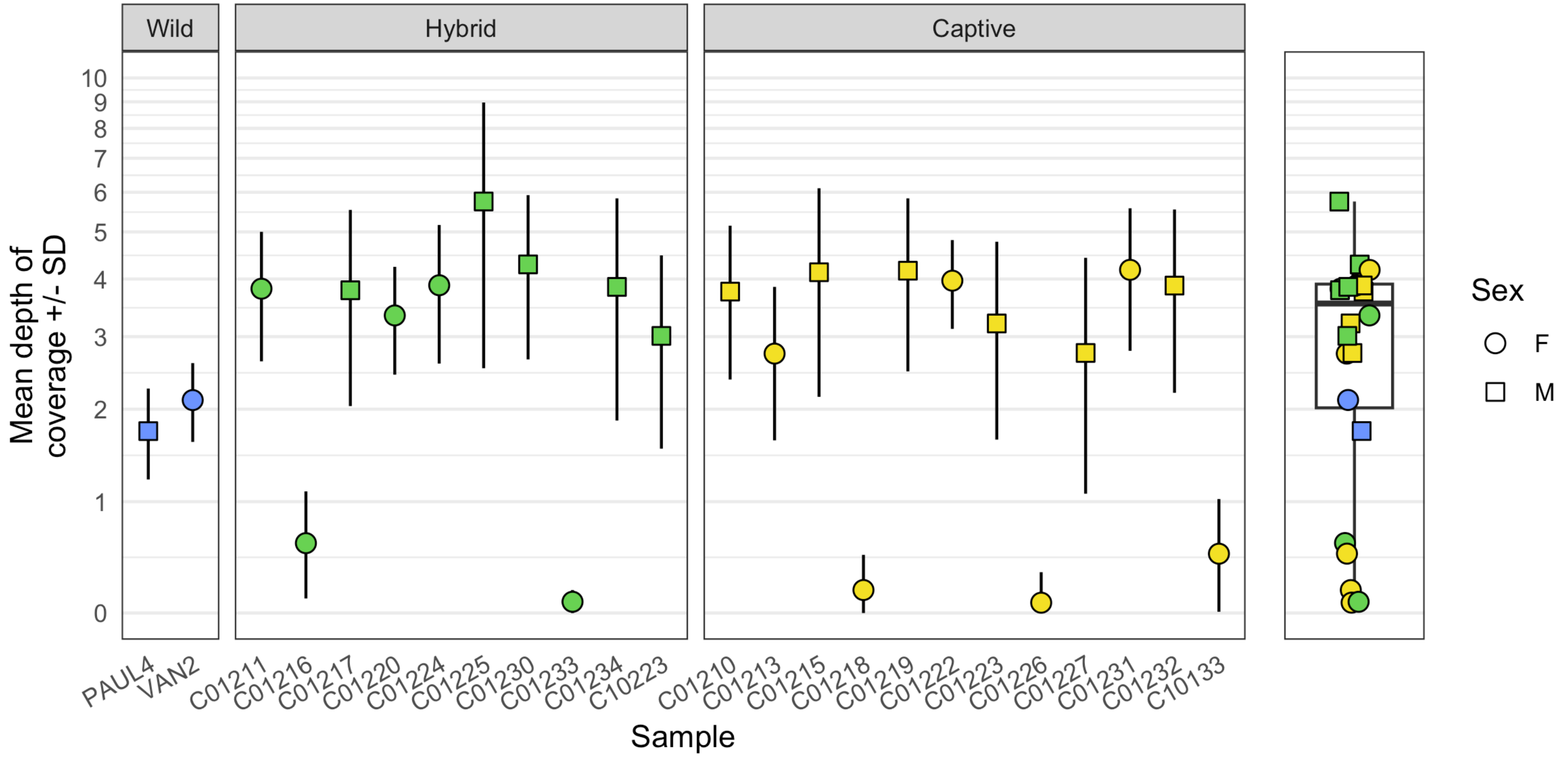


**Supplementary Figure 2.** Sequencing coverage for all individuals across the autosomal scaffolds. Depth was calculated in 100 kbp non-overlapping bins using MosDepth v0.3.1 (Pedersen and Quinlan, 2018) and for each individual means and standard deviations were calculated across all bins. Shapes, which indicate the sex of each individual, represent means and error bars represent ± 1 standard deviation. Where the lower boundary of this range is < 0, the range has been cut off at 0. A boxplot is also shown to help visualise the distribution of mean depths across all samples.

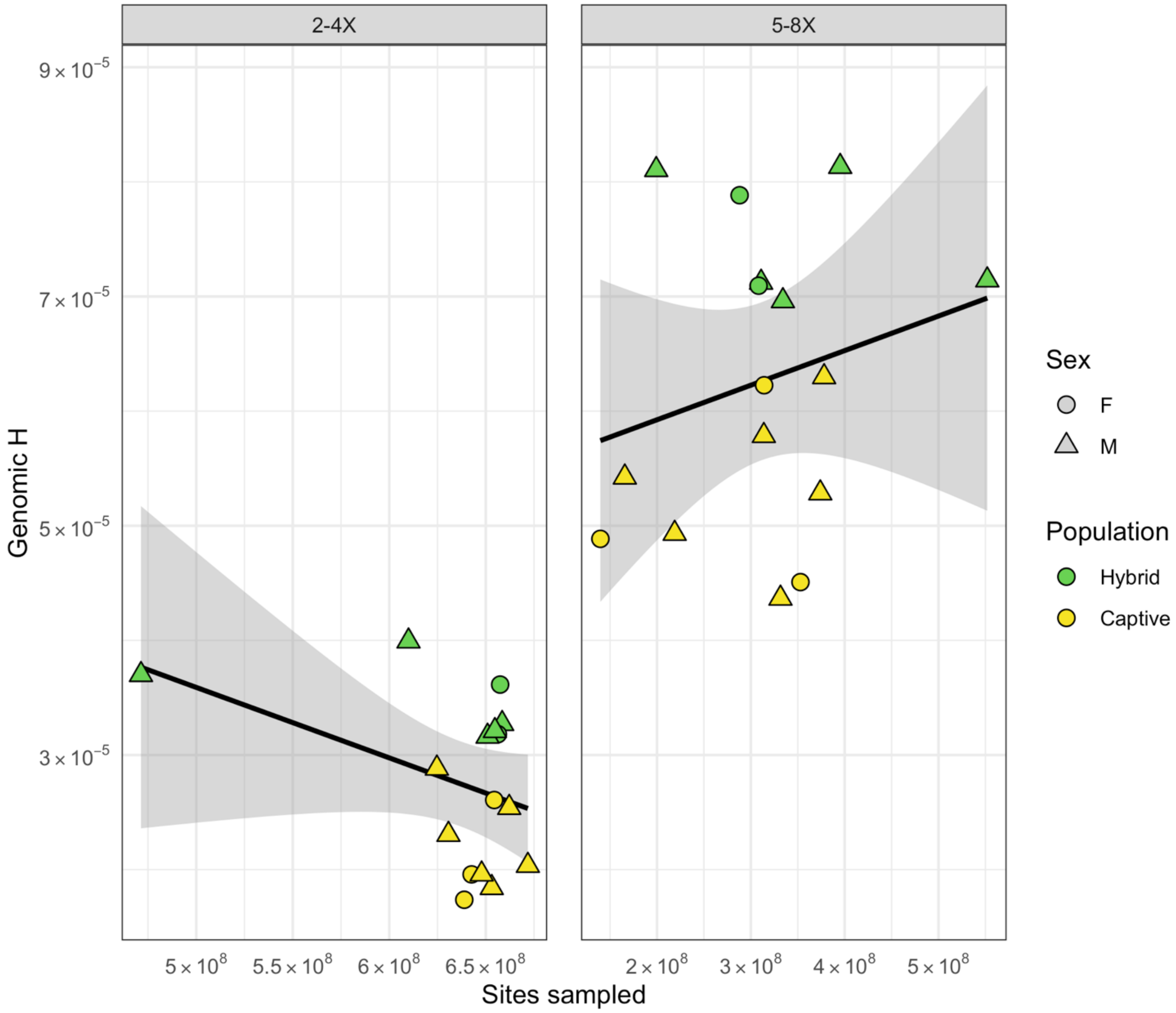


**Supplementary Figure 3.** The relationship between the number of autosomal sites sampled after filtering and heterozygosity, calculated by estimating the single-sample site frequency spectrum for each sample using the -doSaf angsd subroutine. For each individual, the genome was divided into coverage classes of 2-4X and 5-8X. Individuals VAN2, PAUL4, and C01220 (Supplementary Table 1) were excluded. The former two are outliers for both sequencing depth and heterozygosity, and the latter was an outlier for all genetic diversity measures (Figure 3bc) and is likely the result of parthenogenetic reproduction. Black lines and shaded areas represent linear lines of best fit for standard error of the mean estimated by the ggplot2 geom_smooth function. Left: relationship between sampled sites and heterozygosity for the 2-4X site class. Estimated slope = -6.15e-14, $t$ = -1.61, P = 0.13. Right: relationship between sampled sites and heterozygosity for the 5-8X site class. Estimated slope = 3.02e-14, $t$ = -0.91, P = 0.38.

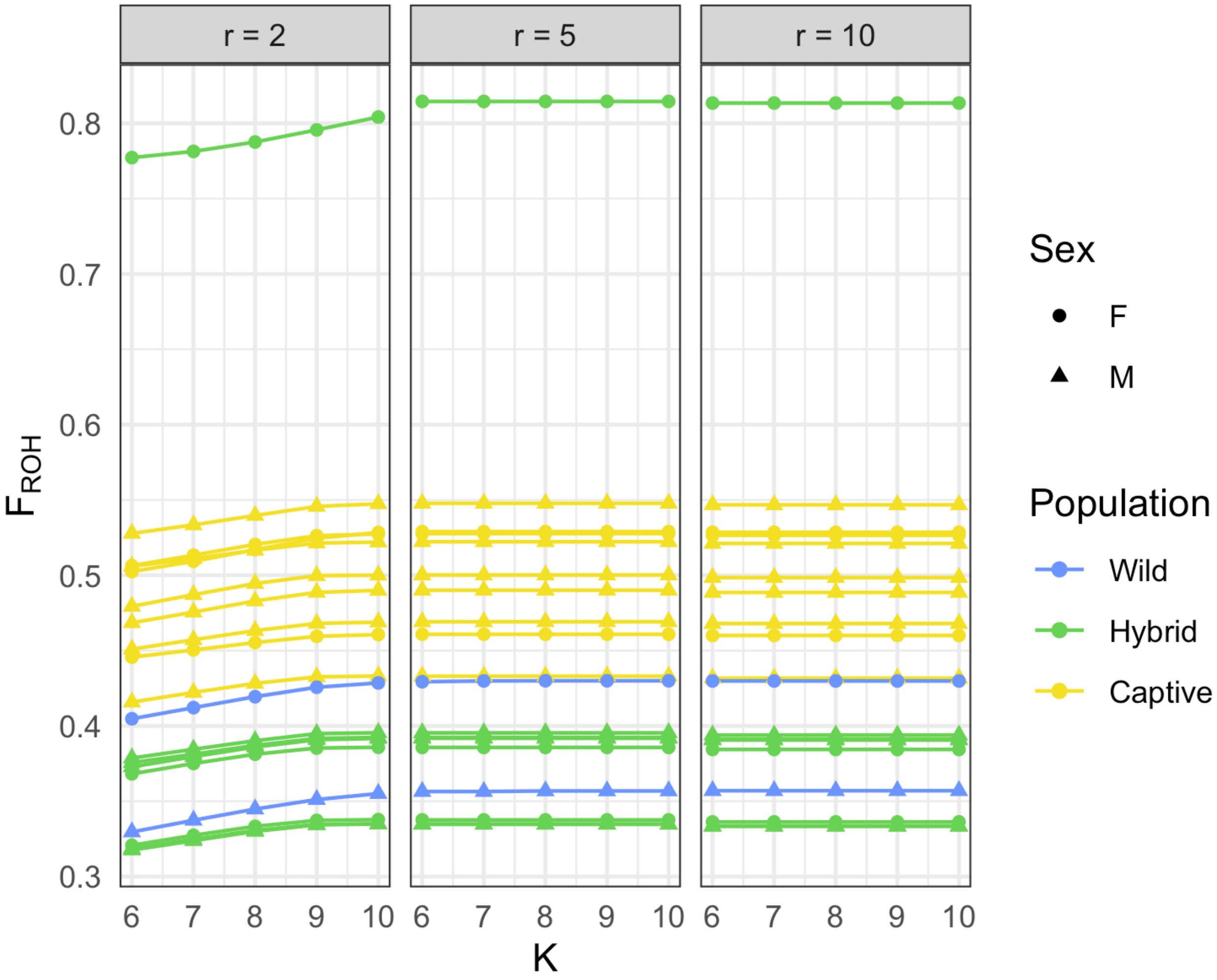


**Supplementary Figure 4.** $F_{ROH}$ estimated using RZooRoH (Bertrand et al., 2019) for models of different K and $R_K$ values. All values taken from models using all individuals to estimate allele frequencies.

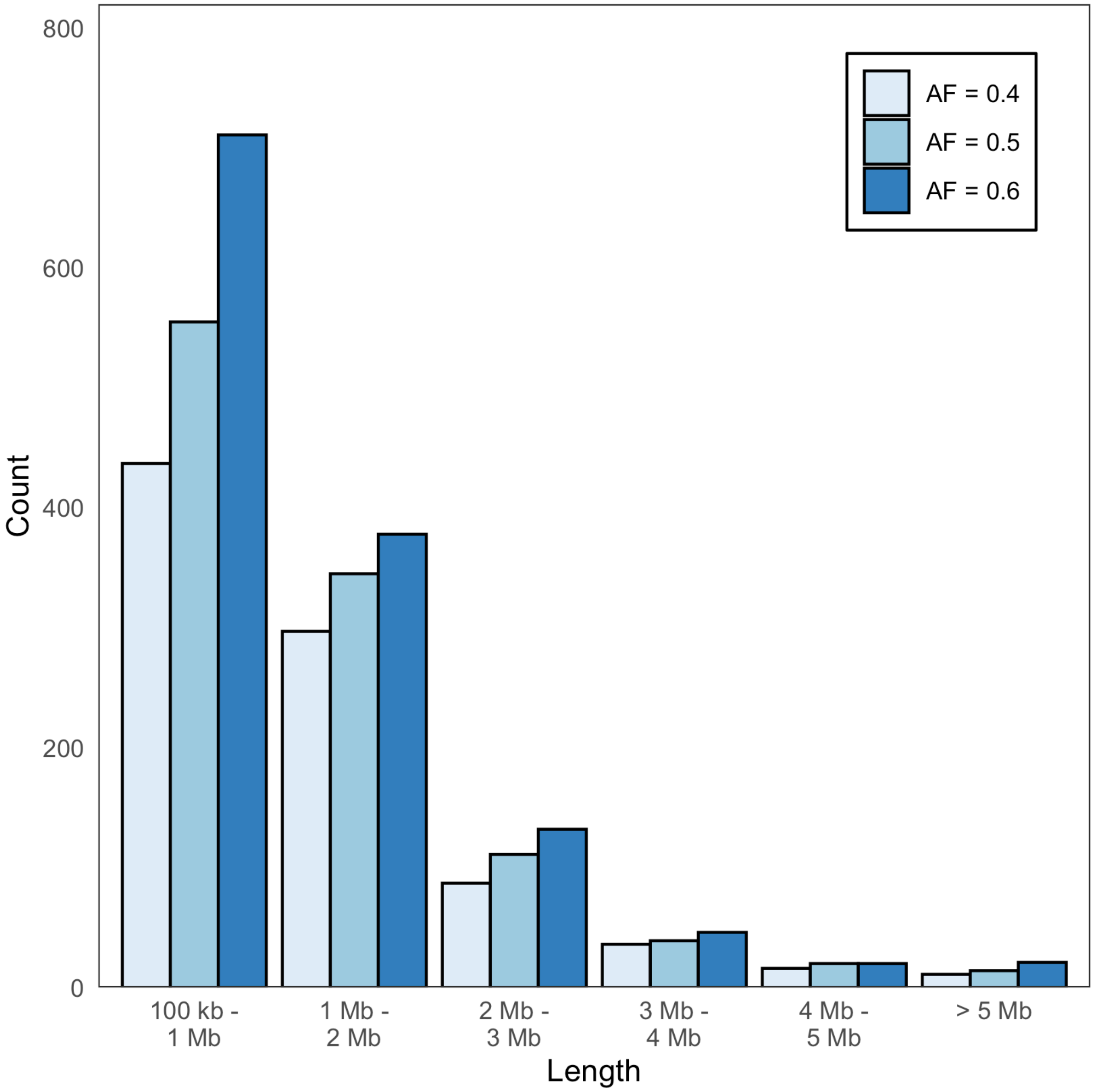


**Supplementary Figure 5.** Summary of ROH detection for a wild sample of *Dryococelus australis*. ROH coordinates were estimated using bcftools v1.19-5-g2bbfdbf9 (Danecek et al., 2021). Multiple starting allele frequency values were used for bcftools indicated by the colour legend. Different minimum length cutoffs are compared for each starting allele frequency.

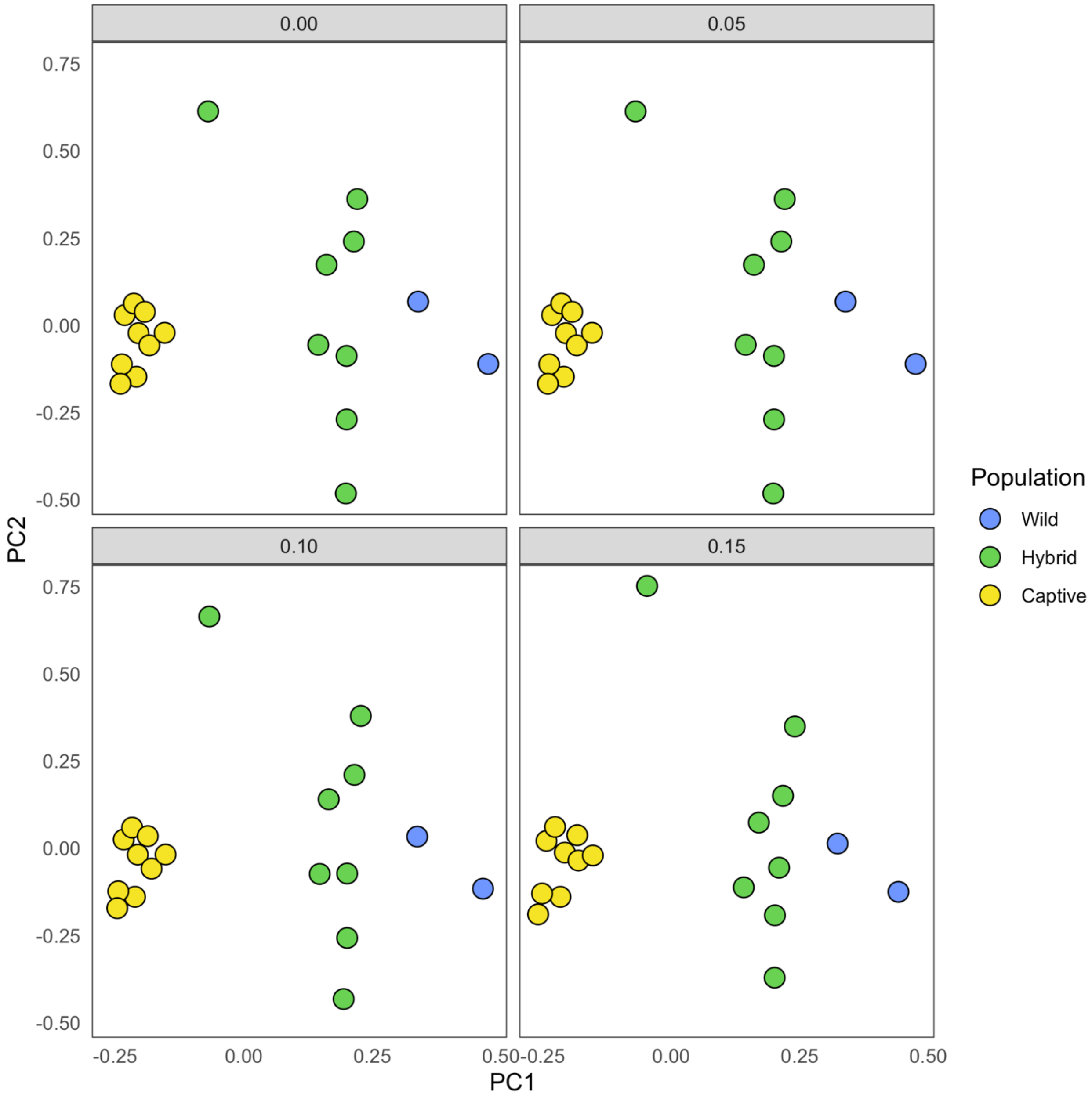


**Supplementary Figure 6.** Population structure among Dryococelus australis samples with different minor allele frequency cutoffs. Covariance matrix estimated using PCangsd v0.98 (Meisner and Albrechtsen, 2018) and eigenvectors calculated in R (R Core Team, 2018).

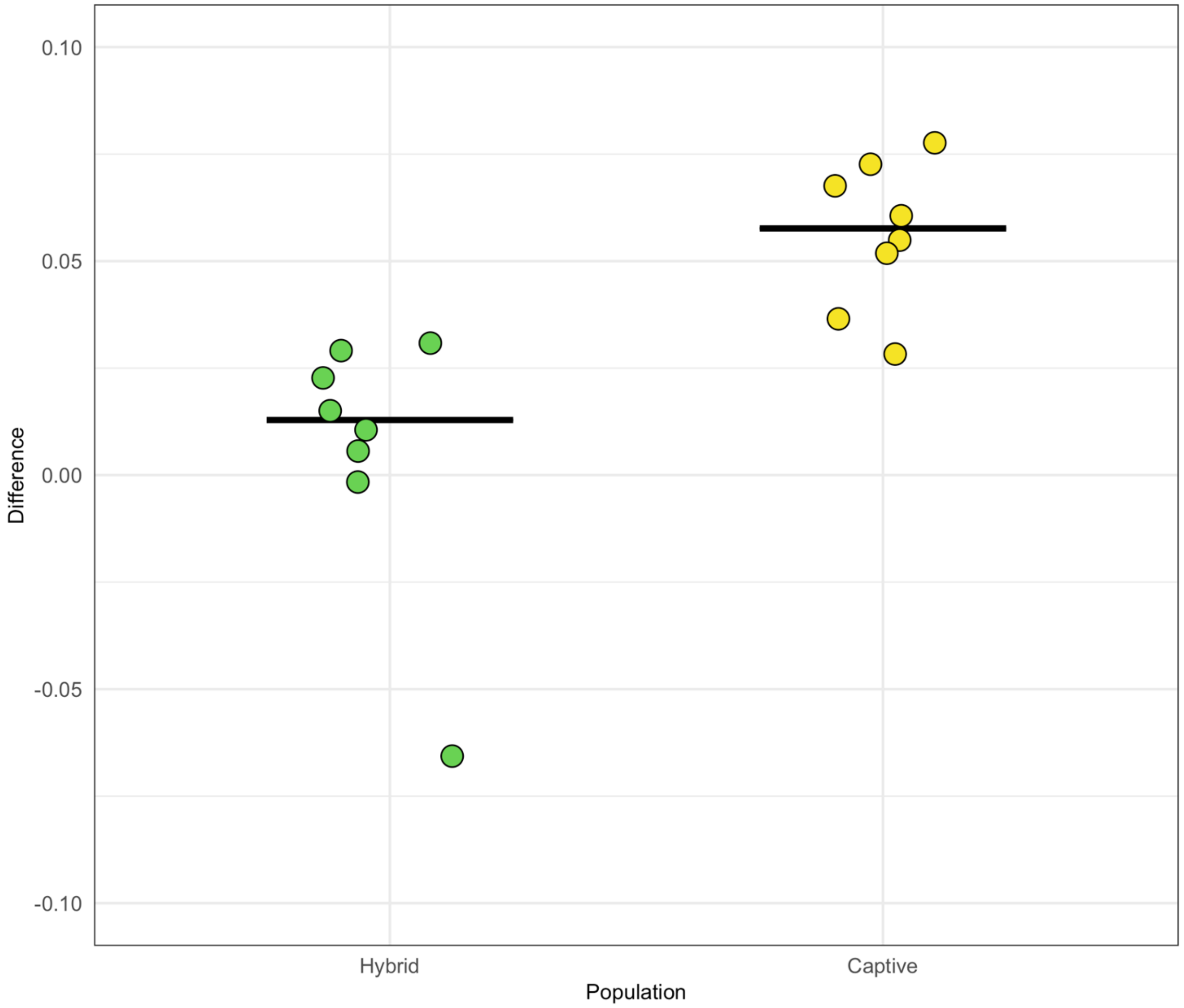


**Supplementary Figure 7.** Difference in $F_{ROH}$ between models including all individuals for allele frequency estimation and group specific models, estimated using RZooRoH (Bertrand et al., 2019). The presented model used values of K and $R_K$ of 9 and 2.

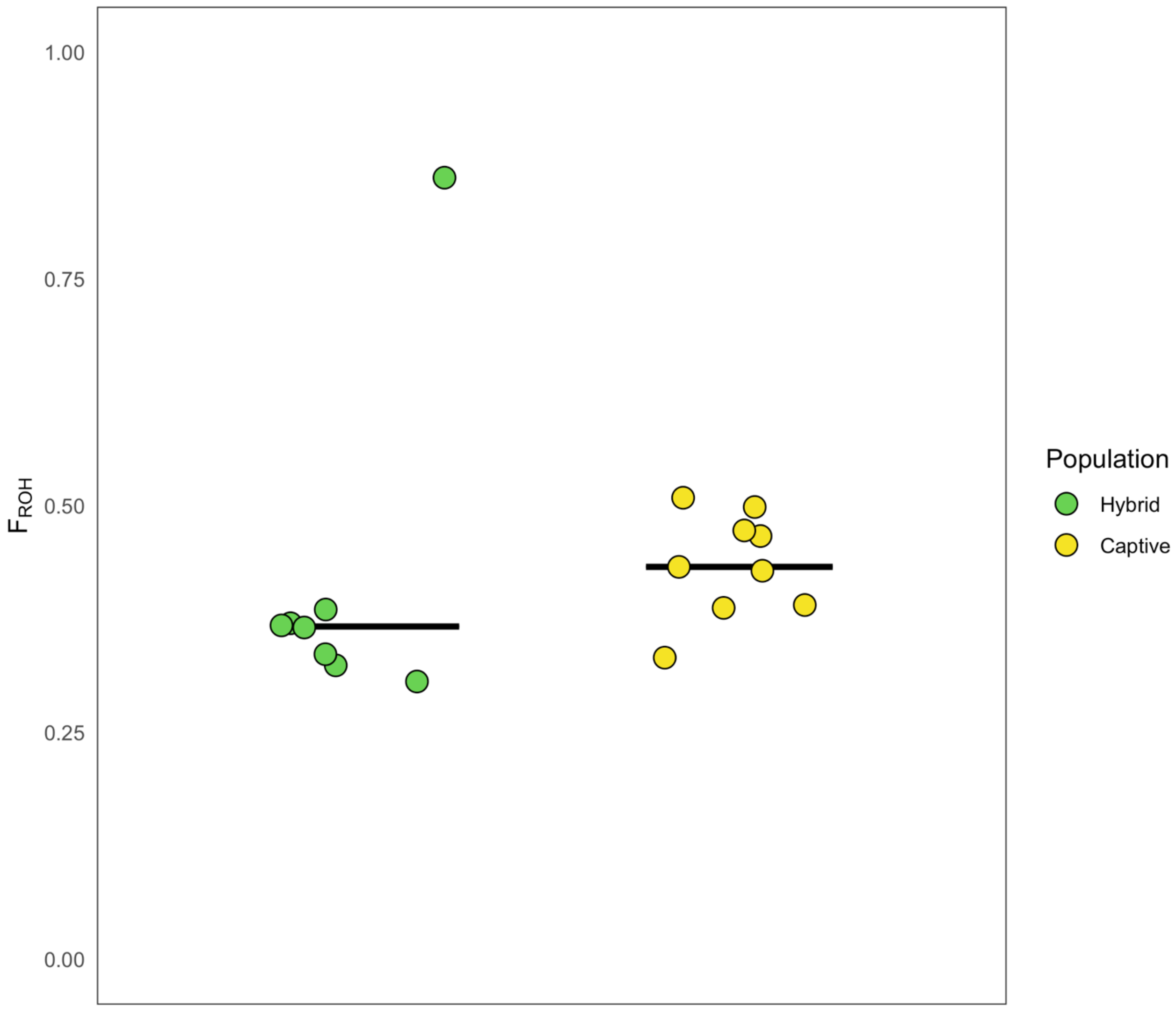


**Supplementary Figure 8.** $F_{ROH}$ in the captive and hybrid groups using population-specific allele frequencies for estimation using RZooRoH (Bertrand et al., 2019). The presented model used values of K and $R_K$ of 9 and 2.

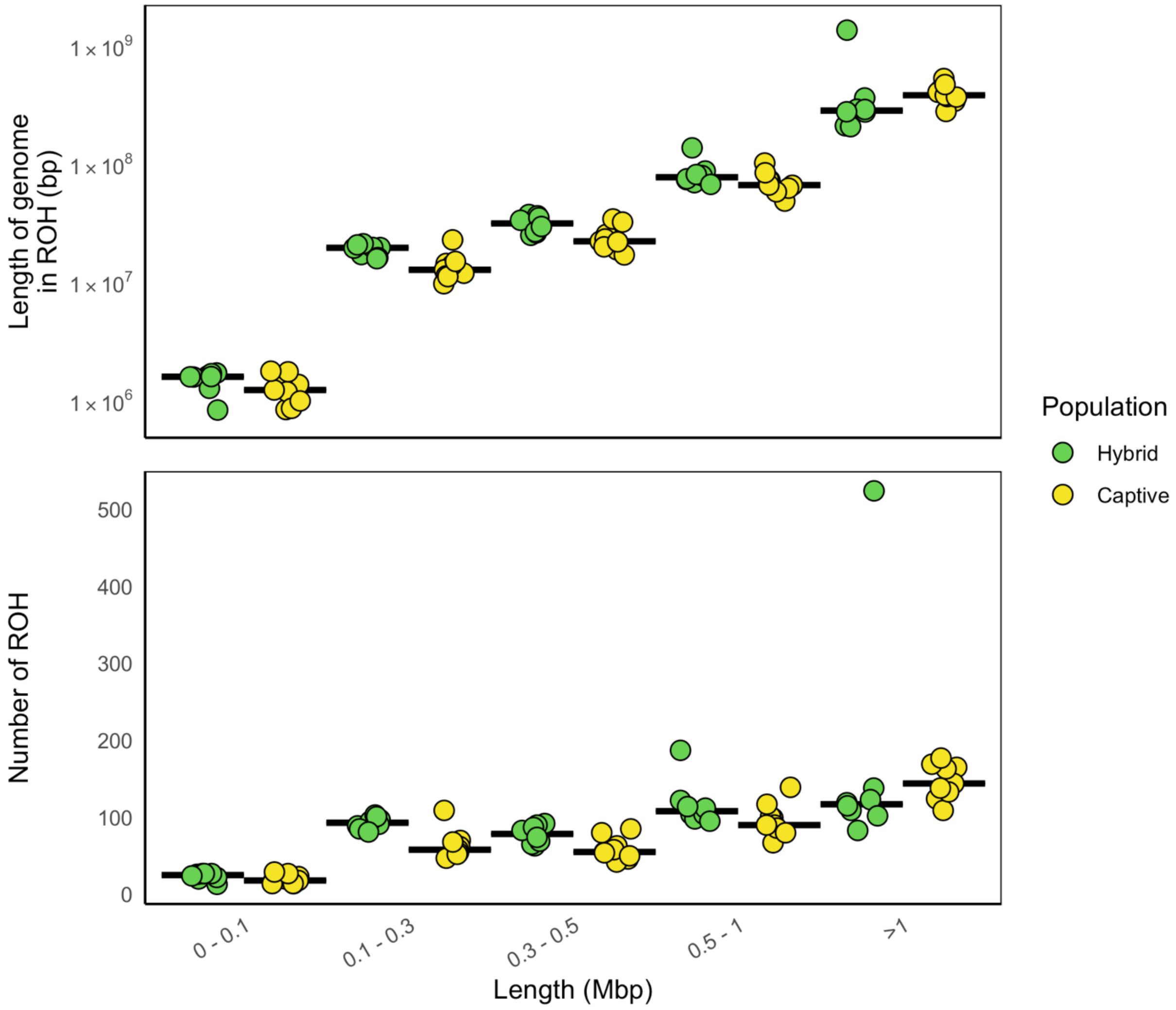


**Supplementary Figure 9.** Summary of runs-of-homozygosity (ROH) detection with RZooRoH package (Bertrand et al., 2019). The models presented here used population specific allele frequencies and K and $R_K$ values of 9 and 2. (left) Cumulative length and (right) total number of ROH per individual across five length classes with black lines group medians. The outlier to the hybrid group in the ‘> 1 Mbp” length class is individual C01220.
